# Course-based Undergraduate Research Module for Enzyme Discovery Using Protein Structure Prediction

**DOI:** 10.1101/2021.08.19.456875

**Authors:** Jessica I. Kelz, Gemma R. Takahashi, Fatemeh Safizadeh, Vesta Farahmand, Marquise G. Crosby, Jose Luis Uribe, Suhn H. Kim, Marc A. Sprague-Piercy, Elizabeth M. Diessner, Brenna Norton-Baker, Steven M. Damo, Rachel W. Martin, Pavan Kadandale

**Author notes:** These authors contributed equally (Jessica I. Kelz, Gemma R. Takahashi).

## Abstract

A major challenge for science educators is teaching foundational concepts while introducing their students to current research. Here we describe an active learning module developed to teach protein structure fundamentals while supporting ongoing research in enzyme discovery. It can be readily implemented in both entry-level and upper-division college biochemistry or biophysics courses. Pre-activity lectures introduced fundamentals of protein secondary structure and provided context for the research projects, while a homework assignment familiarized students with 3D visualization of biomolecules using UCSF Chimera, a free protein structure viewer. The activity is an online survey in which students compare structure elements in papain, a well-characterized cysteine protease from *Carica papaya*, to novel homologous proteases identified from the genomes of an extremophilic microbe (*Halanaerobium praevalens*) and two carnivorous plants (*Drosera capensis* and *Cephalotus follicularis*). Students were then able to identify, with varying levels of accuracy, a number of structural features in cysteine proteases that could expedite the identification of novel or biochemically interesting cysteine proteases for experimental validation in a university laboratory. Student responses to a post-activity survey were largely positive and constructive, describing points in the activity that could be improved and indicating that the activity helped them learn about protein structure.

## INTRODUCTION

The Protein Data Bank (1) contains more than 174,000 structures of biomolecules as of early 2021, and familiarity with protein structures is necessary for understanding the literature in many subfields of biology. Experimentally, protein structures are generally solved using X-ray crystallography, NMR spectroscopy, cryo-EM, or, for complex molecular assemblies, a combination thereof. Advances in experimental methodology, including automated data collection at synchrotron beamlines, improved NMR instrumentation, and the “resolution revolution” in cryo-EM have greatly accelerated the pace of protein structure determination studies. As this methodology becomes easier to use, familiarity with protein structures has become an essential competency needed for many types of biological research. Being able to visualize the relevant molecular structures improves mechanistic understanding of enzyme activity, protein-protein interactions, and regulation of biological processes such as transcription and translation. Connecting protein structure to function has been identified by the American Society for Biochemistry and Molecular Biology (ASBMB) as one of five foundational concepts in molecular biology education, and learning how to relate the primary sequence to three-dimensional structure is a prerequisite for the associated learning goals (2).

Learning to interpret protein structures is therefore one of the fundamental tasks of a student in an introductory biochemistry course. This topic is traditionally considered difficult, and analysis of semantic distance between fields shows that molecular biology and biochemistry are culturally isolated from other disciplines (3). This means a large corpus of field-specific language must be learned starting in the introductory classes, even without considering the information-packed graphical symbology used to express chemical structures. Examples in textbooks and lectures, not to mention the current literature, interchangeably switch between different representations of the same molecules depending on the features being emphasized. Representations where all atoms are shown are generally eschewed because the distracting level of atomic detail obscures the overall fold and key structural motifs, and makes it difficult to locate functional residues without prior knowledge. Space-filling models are useful for building intuition about molecular shape, and with appropriate color coding, surface properties such as charge and hydrophobicity, but they do not allow visualization of the protein interior.

Ribbon or licorice diagrams that omit side chains and individual atoms, and represent *α*-helical and *β*-strand secondary structure elements as coiled helices or flat ribbons, respectively, highlight the three-dimensional organization of the protein. These diagrams were first systematized by Jane Richardson in 1981 (4), although similar drawings had already appeared in individual structural biology papers. Although every introductory biochemistry textbook has a concise explanation of these diagrams, we recommend Richardson’s original review to students who are interested in structural biology: various structural motifs are clearly explained, numerous instructive examples of structural motifs are presented, and the beautiful hand-drawn diagrams highlight the human effort that went into developing this highly efficient representation scheme. Computer programs for automating the production of ribbon diagrams soon followed (5, 6), and modern PDB structure viewers, such as UCSF Chimera (7), PyMol (8), and Visual Molecular Dynamics (9), use these representations as one of the standard settings. Several such programs are available online for free, and are relatively easy to install and use. Here we take advantage of these tools to have students apply their recently gained knowledge about protein structure to an enzyme discovery project using structures predicted from genomic data.

This activity is linked to an ongoing project in the Martin lab, where a major research goal is the discovery of novel enzymes from genome and transcriptome data, in particular from carnivorous plants. These plants have adapted to grow in nutrient-poor environments by obtaining much of their nitrogen from protein in insect prey (10). Carnivorous plants are expected to have a variety of proteases with different activities, as they rely on these enzymes for digestion as well as the more typical functions of plant proteases: cellular housekeeping, defense against insects and pathogens, and hydrolysis of seed storage proteins. In the Venus flytrap (*Dionaea muscipula*), expression of at least one digestive protease is upregulated in response to prey stimuli (11). As expected, the genomes of the Cape sundew (*Drosera capensis*) (12) and the Albany pitcher plant (*Cephalotus follicularis*) (13) have yielded many new proteases — so many, that the main problem is choosing appropriate targets for experimental investigation. In general, determination of experimental structures is a bottleneck for enzyme discovery from nucleic acid sequencing data. Advances in sequencing methodology have outstripped even the rapid pace of development in structural biology methods, in part because of the difficulties inherent to sample preparation. Preparing protein samples of sufficient quantity and purity for structural studies is time-consuming and expensive, and requires extensive training and experience, as does interpretation of the data. Performing these experiments is impractical for every putative enzyme discovered from a genome or transcriptome. Therefore, we use structural models derived from sequence data using protein structure prediction tools such as Rosetta (14, 15) and iTasser (16). Although the predicted structures do not capture every detail, particularly when considering side chain conformations, we find that they are highly reliable for predicting the overall folds of enzymes belonging to well-known structural classes, including the cysteine proteases used in this activity. This was illustrated by the crystal structure of a cysteine protease from *D. muscipula* (17), which was solved after we predicted its structure (18). Our predicted structure has excellent overall agreement with the experimental one, and captures all of the functionally important features of the active site. Results such as this as well as ongoing validation efforts such as the CASP competition (19), provide evidence that structures predicted in this manner are sufficient to verify functional folds and active sites for well-known enzyme classes. With recent machine learning-based advances in protein structure prediction such as AlphaFold (20) and RoseTTAFold (21), it is now possible to obtain large numbers of predicted structures for members of an enzyme class of interest, such that the activity can be updated frequently or tailored to fit the theme of a particular class.

Predicting structures *en masse* for enzymes discovered from genomic data provides a foundation for predicting which proteins will have functional differences from well-characterized members of the same enzyme class; however, examination of the structures and prediction of functionality is not easily automated. Some features, such as extra domains, are apparent from the sequence alone and could be detected using standard software tools. Others are more subtle and require examination by a human with some training in protein structure analysis. For instance, even relatively small occluding loops can dramatically alter substrate specificity by partially blocking the active site cleft, and these cannot necessarily be identified in sequence space, as they interact with the active site cleft in three dimensions. Fortunately, given an appropriate reference protein, undergraduate biochemistry students can learn to identify such features relatively quickly in the context of a class activity. Here we describe such a course-based undergraduate research experience (CURE) module for students in an undergraduate biochemistry class. Students received training in protein sequence and structure analysis, and then worked individually to identify similarities and differences between papain, a well-characterized plant cysteine protease, and a novel protein from either *D. capensis*, *C. follicularis*, or the extremophilic microbe, *Halanaerobium praevalens* (22).

## SCIENTIFIC AND PEDAGOGIC BACKGROUND

A major challenge in teaching protein structure interpretation is that the connection between the intermolecular forces holding proteins together and the three-dimensional structures that result is abstract. Furthermore, many students enter introductory biochemistry with limited three-dimensional visualization skills, such that practicing a task that requires manipulating protein structures in a virtual 3D environment is helpful. The examples presented in introductory textbooks are often selected to present a wide range of different structural motifs, which provides a good overview of existing structures, but can come across as disconnected. Here we introduce a particular enzyme class, cysteine proteases (MEROPS family C1) (23), and invite students to look for relatively subtle structural differences. We selected cysteine proteases because there are a large number of characterized structures for this enzyme class in the PDB, making structure prediction very useful for determining overall folds and relative domain orientations. At the same time, there are no shortage of newly discovered and uncharacterized cysteine proteases, as many plants have multiple paralogs of these common defensive proteins (24, 25), of which only a few have been studied in detail. *D. capensis* has 44 cysteine proteases (18), which we have previously modeled and categorized according to the classification scheme of Richau et al. (24), while *C. follicularis* has at least 16 (13). Our protein set consisted of the 16 novel papain-like cysteine proteases from *C. follicularis*, matched with 17 cysteine proteases from *D. capensis*, whose structural features had already been examined by the Martin group. One additional cysteine protease from the extremophilic microbe, *H. praevalens*, was also included in order to assess the robustness of this characterization method when examining proteins that are less closely related. Each student was assigned a unique protease from this set of 34, and all students used the crystal structure of papain from *Carica papaya* (UniProt ID: PAPA1 CARPA; PDB ID: 9PAP; (26)) as a reference protein to compare structural features. The main objectives of this class activity were to introduce students to the basics of protein structure, to help them examine and manipulate protein structures in a virtual 3D environment, and to provide an opportunity to participate in a live research project focused on enzyme discovery.

Course-based Undergraduate Research Experiences (CUREs) have numerous benefits for students, including making research experiences more equitablly available to all students (27), increasing scientific affect (28), improving scientific skills (29), and increasing student retention (30). Further, participation in CUREs early in their university experience improved the odds of students graduating with a STEM degree, and improved student GPAs when they graduated (31). Shorter-term gains from CUREs included improved content knowledge, increased prob-ablity of pursuing longer-term, apprenticeship-based research experiences prior to graduation (30, 32) and abrogation of some so called “achievement gaps” for minoritized students (33). Traditionally, CUREs have been implemented either in lab courses, or in the lab sections of theory courses. To the best of our knowledge, no one has implemented a CURE module within a regular, large lecture course. Given the numerous benefits of exposing students to research experiences, we sought to create a research experience embedded within a lecture course that was based on our enzyme discovery research.

As a pilot for the large class, we first performed the activity via Zoom with undergraduate students in Chem341L (Physical Chemistry Lab) an upper-division course at Fisk University, a private historically black university in Nashville, Tennessee (October 2020). There is precedent for sophisticated protein structure activities in upper-division biophysical courses such as this. For example, undergraduate students assigned to solve the crystal structure of a small protein from its electron density map were very successful even without knowledge of the protein sequence, modeling ambiguous residues using chemical knowledge to identify local interactions, and in some cases producing a better result than the original structure (34). Other activities have focused on using molecular dynamics tools to teach structure visualization, ligand interactions (35), and noncovalent inter-residue interactions (36).

In this activity, graduate students taught a lesson introducing general protein structure concepts and important structural features of proteases in particular. The lecture material focused on protein secondary and tertiary protein structure, with examples of types of secondary structures found in globular proteins as well as the importance of intrinsically disordered proteins. There was also an informal and highly interactive class discussion around current protease projects in the Martin lab, including the carnivorous plant proteins in this dataset, as well as the SARS-CoV-2 main protease (M*^pro^*), which served as a transition into the hands-on activity. The goal of activity was to help students solidify their knowledge and exercise what they learned from the lecture, using their new insight to help discover novel structural features in papain-like protein structures. Due to the small class size (9 students) and the students’ relatively advanced knowledge of molecular structure, each student was able to examine multiple structures and compare notes about different protein features, including pro-sequences, granulin domains, and differing degrees of active site cohesion. 3D-printed structures of selected proteins were provided, as there is evidence that examining 3D-printed models of protein structures helps students build accurate mental models of protein structure (37).

In order to incorporate this module into a large lecture course, we created a shorter version that we implemented in two sections of a lower division Biochemistry course. This was a large enrollment class (356 students in one section, and 252 students in the other section), required for all students in several majors, including Biology, Pharmaceutical Sciences, Nursing, and Public Health. The course is taught as a one-quarter survey course of major concepts in Biochemistry, including amino acid properties, and protein structure and function.

In the rest of this paper, we describe the design of lecture materials and the cysteine protease survey, and discuss the results of the activity and its assessment, which we hope will be useful for other biochemistry educators. The survey materials and the protein models used are provided in the Supplementary Information.

## MATERIALS AND METHODS

### Protein sequences and structural models

Sequence alignments were performed using ClustalOmega (38), with settings for gap open penalty = 10.0 and gap extension penalty = 0.05, hydrophilic residues = GPSNDQERK, and the BLOSUM weight matrix. For the *D. capensis* proteases, the presence and position of a signal sequence flagging the protein for secretion was predicted using SignalP 4.1 (39). An initial model was created for each complete sequence using the Robetta (14) implementation of Rosetta (15). Any residues not present in the mature protein were removed, disulfide bonds identified by homology to papain were added, and the protonation states of active site residues were fixed to their literature values. Each corrected structure was then equilibrated in explicit solvent under periodic boundary conditions in NAMD (40) using the CHARMM22 forcefield (41) with the CMAP correction (42) and the TIP3P model for water (43); following this minimization, each structure was simulated at 293K for 500ps, with the final conformation retained for subsequent analysis. The published structure of papain (PDB ID: 9PAP; (26)) was used as the initial starting model (following removal of heteroatoms and protonation using REDUCE (44)), and similarly equilibrated prior to use as a reference.

For the proteases from *C. follicularis* and *H. praevalens*, retrieved from UniProt (45) (Table S1), structure prediction was performed using iTasser (16). Signal sequences were not removed from these proteins in order to leave them as a point of discussion for the class activity.

### The cysteine protease survey

The cysteine protease survey was designed to guide students through the process of comparing a novel cysteine protease structure to that of papain in UCSF Chimera. Questions identified characteristics like various secondary structure locations, blocked active sites, and relative lengths of N- and C-termini. The full survey can be found in the Supplementary Information.

### Post-activity survey

After completion of the activity, students were asked to answer a questionnaire about their experience. The survey was administered in Canvas as a regular weekly activity for the class. The questions were: 1. In how many classes at UCI (prior to this one) did you have the opportunity to apply the concepts you were learning about in class to a research project? 2. Please tell us what you liked best about the project. 3. Please tell us what you liked least about the project. 4. Do you agree or disagree with the following statement: This research project helped me understand protein structure-function better. 5. Do you agree or disagree with the following statement: This research project should continue to be a part of this course. and 6. How can this research project be improved?

## RESULTS AND DISCUSSION

### Pre-activity training

During the class before the protease discovery activity, a general introduction to protein structure was presented. The concepts of primary, secondary, and tertiary structures were presented, along with a primer on interpreting ribbon diagrams. Examples are shown in Figure 1.

**Figure 1:**
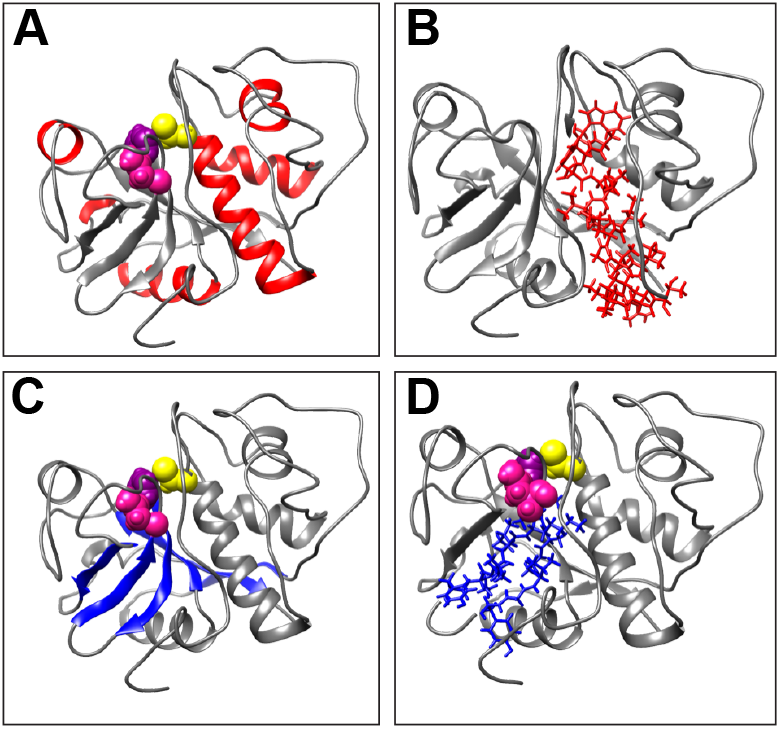
Papain secondary structure examples presented in pre-survey lecture. A) All *α*-helices (red) displayed as ribbons. B) One *α*-helix (red) displayed with all atoms shown as stick models. C) All *β*-strands (blue) displayed as ribbons. D) Two *β*-strands (blue) displayed with all atoms shown as stick models.

Before starting the in-class exercise, an introductory lecture on cysteine protease discovery was presented, taking approximately 20 minutes. This lecture was delivered by a graduate student directly involved in the research, and began by describing the motivation for discovering new cysteine proteases. Examples presented include finding highly specific proteases to cleave expression tags or break down proteins into smaller peptides for bottom-up proteomics, and on the other hand finding very general proteases to break down biofilms and cleave protease-resistant aggregates such as amyloid fibrils. The overall workflow of the project was summarized, emphasizing the large number of proteases discovered from the *D. capensis* genome and how molecular modeling could help narrow down the targets chosen for experimental characterization. Finally, some examples of *D. capensis* cysteine proteases with functional features different from papain were shown. The example proteases are shown in Figure 2. The first, aspain, has an unusual active site configuration with an aspartic acid taking the place of the canonical asparagine and a large occluding loop partially blocking the active site, potentially modulating substrate specificity. The second, DCAP _6097, has a C-terminal granulin domain, which is common in proteases that cleave storage proteins during seed sprouting. Both contain examples of structures students may encounter when studying novel papain-like proteases.

**Figure 2:**
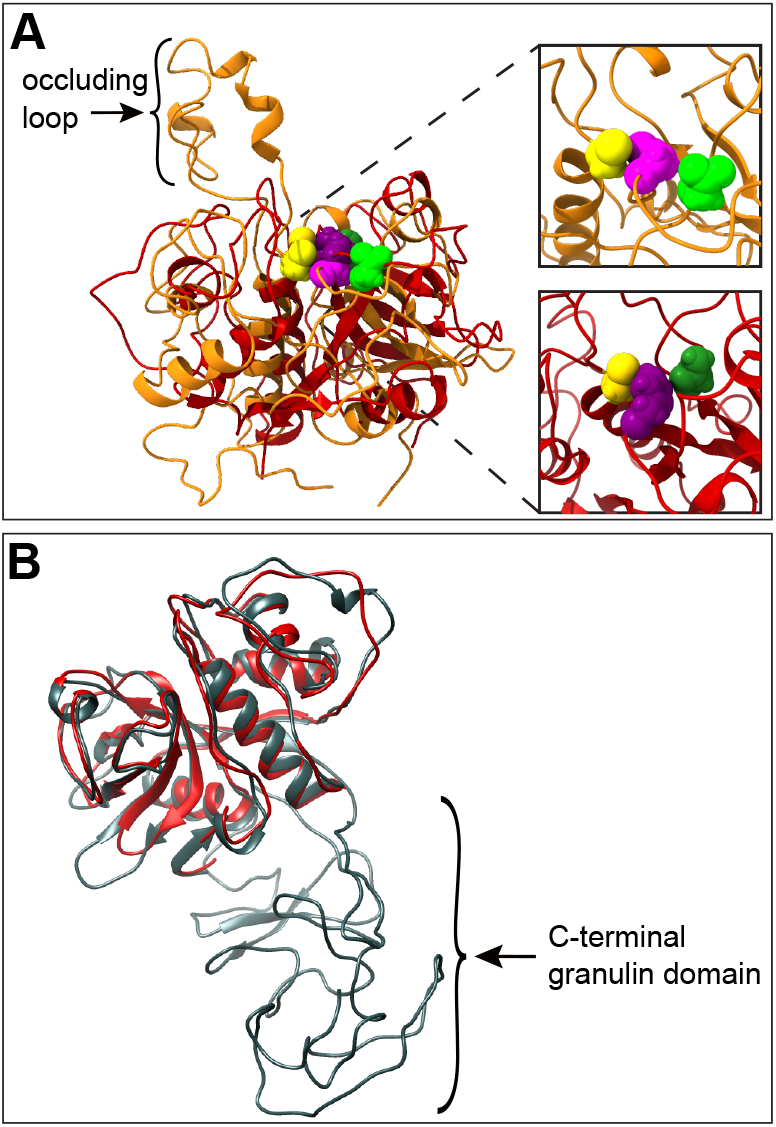
Example cysteine proteases, aligned with papain (red), presented to students prior to taking the in-class survey. A) Aspain: DCAP _3968 (orange). Aspain’s unusual active site (top inset) replaces the typical asparagine (dark green) of papain (bottom inset) with aspartic acid (lime green). Its occluding loop, which partially blocks the active site, is indicated by an arrow. Other active site residues: cysteine: gold/yellow; histidine: purple/magenta for papain and aspain, respectively. B) DCAP _6097 (dark grey). DCAP _6097’s C-terminal granulin domain, indicated by an arrow, extends well beyond the rest of the papain-aligned structure.

### In-class exercise

In order to provide practical experience with comparing structurally related proteins, we assigned each student a protein from our set of predicted structures, which they were instructed to compare to papain. Every student was given two .pdb files to download, the reference papain structure and the “new” structure. An example is shown in Figure 3. The structures for papain (Figure 3A) and the novel protein DCAP 4793 (Figure 3B) are very similar in overall fold, and differences are difficult to determine when examining them separately. However, overlaying the structures (Figure 3C) reveals some potentially functionally relevant differences. The region labeled 1 shows the difference in length of two *β*-strands and the loop connecting them: both the strands and the loop are longer in DCAP _4793 than in papain. The area labeled 2 shows a short *α*-helix in DCAP _4793 that is absent in papain. Both proteins have a long helix ending in the area labeled 3, but it is longer in papain than in DCAP _4793. Differences in backbone position of the long loops are also observed (e.g. in the region labeled 4), but these are considered to be a result of variable dynamics in these structural elements rather than persistent, meaningful differences. Discussion of which of these structural differences are likely to be functionally relevant was arguably the most difficult part of the exercise, and at the same time led to valuable conversations about the types of judgement calls made by structural biologists and how protein structure can serve as a starting point for hypotheses about function.

**Figure 3:**
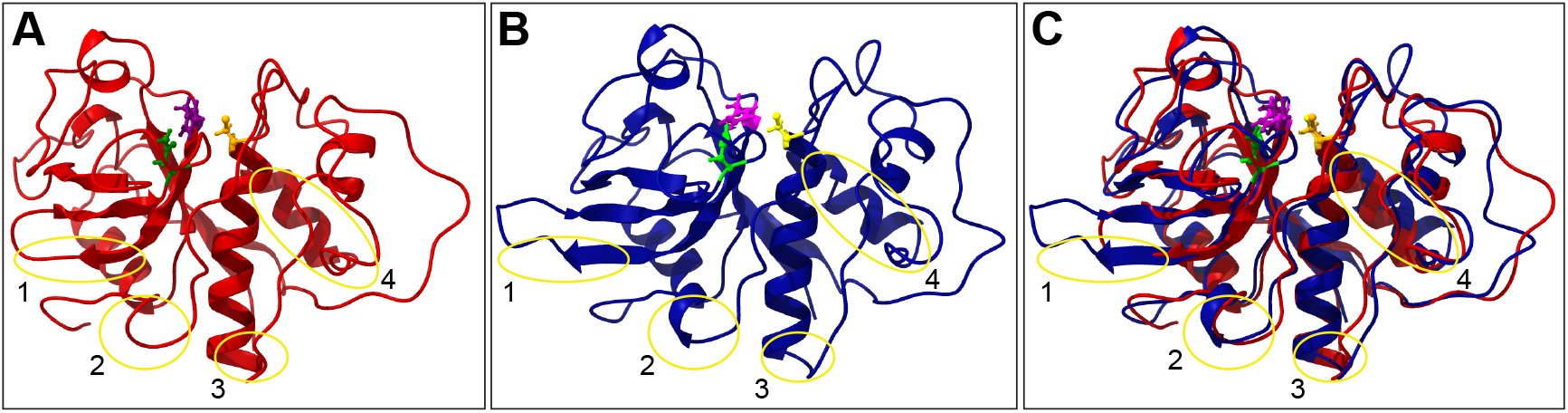
Comparison of the reference papain structure to a molecular model of a new protein, DCAP _4793. A) The papain structure is shown in red. Circled areas (cyan) highlight differences in comparison to DCAP _4793. B) The molecular model for DCAP _4793, generated using Rosetta, is shown in blue. C) An overlay of the two proteins in A and B highlights similarities and differences described in the main text. The active site residues in both proteins are shown as space-filling models with color codes as follows: cysteine (gold/yellow); histidine (purple/magenta); asparagine (dark/lime green) for papain and DCAP 4793, respectively.

### Detection of novel protease features

Not all protease features were interpreted in the same way; some were correctly identified by most participants, while others received mixed responses of varying accuracy. Students did, for example, correctly match most large *α*-helices to those in papain (Figure 4A and D), but often struggled to identify partially or fully blocked active sites (Figure 4C and D). Furthermore, more ambiguous structural features, like papain’s small sixth *α*-helix (Figure 4B and D), were identified with mixed levels of success. Representative data for several of these questions are shown in Figure 4: Q4: Is there an *α*-helix on your structure that lines up with the first *α*-helix in papain? (yes/no); Q5.5: Is there an *α*-helix on your structure that lines up with the sixth *α*-helix in papain? (yes/no); Q14: Do you see a feature that is partially or fully blocking the active site? (yes/no); Q18: What differences does your protein have when compared to papain that were either not fully captured or not addressed at all in earlier questions? (free response). For DCAP _5945 (Figure 4A), Q4 and Q14 were answered accurately, as this protein does have an *α*-helix that matches papain’s first *α*-helix, and does not appear to have a blocked active site. In the free response to Q18, most students also suggested the presence of DCAP _5945’s granulin domain, describing a much longer sequence and extra secondary structure elements. DCAP _5945’s Q18 response bar shows that a number of students responded with some identifying description of this granulin domain (yes they did or no they did not). These responses demonstrate what students did very well: identify large structural features that were clearly explained in pre-survey presentations. Other questions, however, did not receive such consistent answers. Papain’s sixth *α*-helix is an example of a more ambiguous structural feature, whose presence or absence in other proteins is subject to interpretation. For example, DCAP 6547 (Figure 4B) does contain an *α*-helix near papain’s sixth *α*-helix, but a lack of overlapping residues and some variation in local torsion angles make it difficult to judge if these are truly aligned; in this case, both “yes” and “no” are reasonable answers to Q5.5. In addition, most students did not recognize a large N-terminal pro-sequence blocking the active site in many proteins, answering “no” to Q14; this can be seen in the responses given in Figure 4B and D. When viewing the accuracy of student responses as a whole (Figure 4D), clear differences emerge between questions. Q4 was answered with relatively high levels of accuracy, while Q5.5 received responses of mixed accuracy, although there was no unambiguously correct answer for several proteins. In contrast to the largely accurate responses of Q4 and Q5.5, in Q14, students were able to identify active sites that were not blocked with good accuracy, but did have difficulty identifying blocked active sites. This suggests that more instruction should be given on this topic in future implementations of the activity.

**Figure 4:**
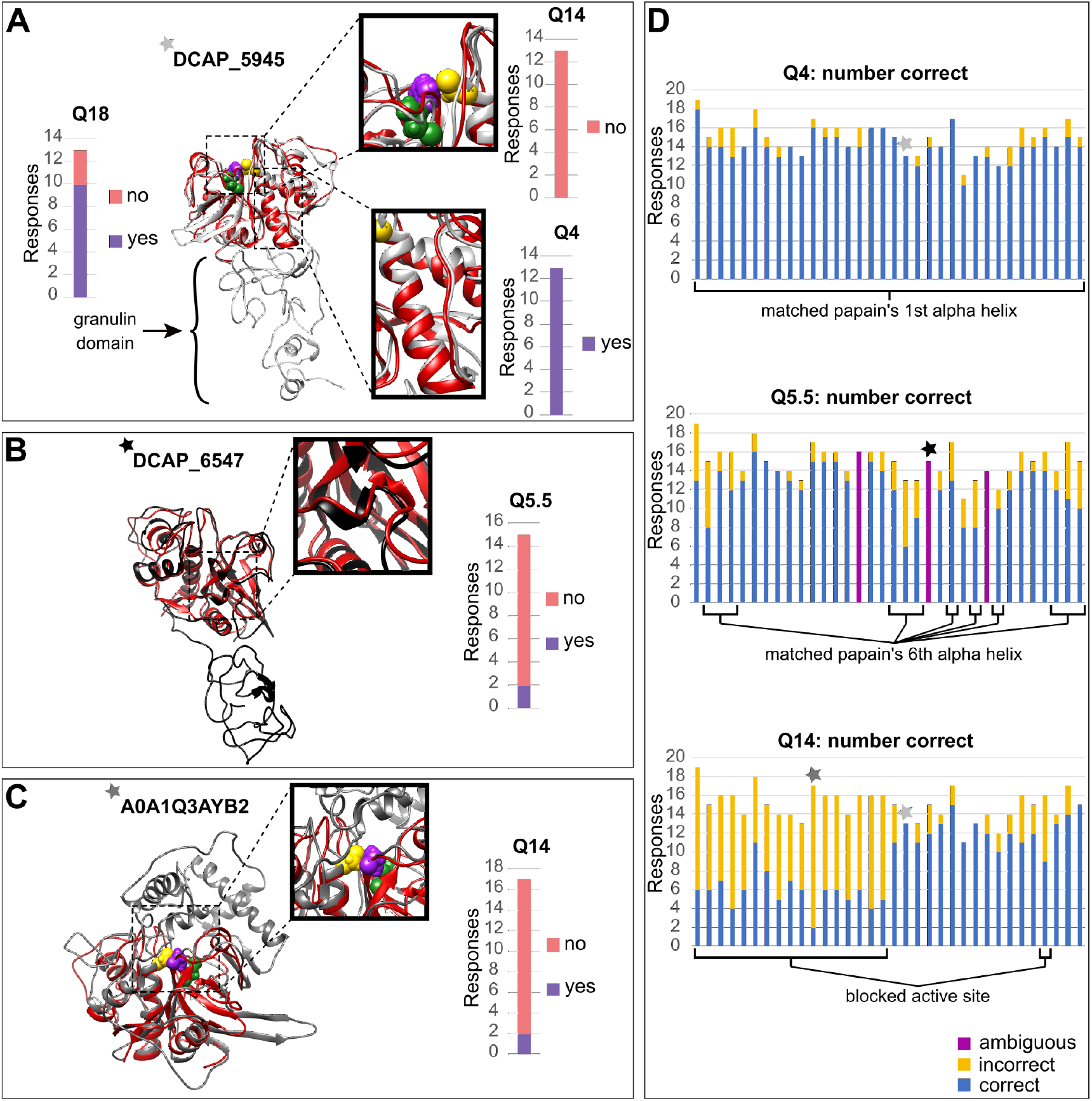
Example survey questions and student responses using proteins aligned to papain (red). A) Examples of accurate and informative student responses using DCAP _5945 (light grey). B) Example of ambiguity in student responses using DCAP _6547 (black). C) Example of inaccuracy in student responses using *C. follicularis* protein A0A1Q3AYB2 (dark grey). D) Accuracy of all student responses to Q4, Q5.5, and Q14. All 34 proteins are shown in each panel, and those whose examples appear in parts A, B, and C are indicated by colored stars (DCAP _5945: light grey; DCAP _6547: black; A0A1Q3AYB2: dark grey). Black brackets below each graph show which subsets of proteins contain the feature in question.

Discrepancies may have a number of causes, including the inherent difficulty of capturing snapshots of certain dynamic protein features (like very short *α*-helices, or flexible termini), differences in survey interpretation, and use of structural cues, rather than Chimera’s predictive software for secondary structure identification. For example, the ambiguous alignment of papain’s sixth *α*-helix in several proteins (Figure 4B and D) is likely a result of the torsion angle cutoff used to define true *α*-helices in Chimera; despite the clear visual alignment of these coil-like structures, part or all of their residues may not be considered *α*-helical in nature. These results speak to the importance of both clarity in what is being asked of participants, as well as emphasis on natural variation of the structural patterns they are asked to characterize. For many of these features, however, different responses are simply a result of varied, but equally valid interpretations of ambiguous data. This kind of harmless variance contributes to the strength of crowdsourced studies, and allows researchers to note potentially mobile or disordered regions. Consequently, future iterations will work to refine the organization and clarity of pre-survey presentations and survey questions, without biasing students’ answers.

### Student experience assessment

Students’ responses to the questions about their experience with the activity are summarized in the tables below. Results are not mutually exclusive, as multiple features were coded from each answer where applicable. Therefore, the number of responses in each category is larger than the total number of respondents. Table 1 shows in how many classes students were given the opportunity to apply concepts learned in class to a research project. Most students had never performed a similar activity in a class before, although some reported as many as three such experiences. Table 2 summarizes the most common responses given for what students liked best about the project. The most common responses cited the interactivity of the activity, seeing how concepts learned in class applied to real-world examples, and having the opportunity to contribute to an ongoing research project. Many students mentioned applying their knowledge to a real-world problem (93 responses) or knowing their work would contribute to an active research project (90 responses), e.g. “I really enjoyed putting what I have learned to use! It really motivated me to work hard on that assignment and to pay attention in lecture, as I knew it had pertinent information I would need.” Others focused on the interactive format of the exercise (82 responses) and the ability to view the proteins in 3D, examine them from different angles, and correlate sequence with structural features (86 responses), none of which are possible with a picture in a textbook. “What I like the most about this project is that I got to look at the protein in 3D, and it is very interesting. On the textbook or online, the protein are always 2D and we cannot spin it around to see its structure.” Some students specifically stated that doing the activity helped them understand protein features (59 responses), and others indicated that it was fun (30 responses). Fifty-five students cited using the UCSF Chimera software as one of their favorite aspects of the project, with several of them explaining that they enjoyed learning a tool that is used by researchers working on protein structures. “I loved the program Chimera and how easy it was to visualize the protein. It was very interesting to compare the different structures to each other based on their sequencing. I felt like a real scientist” and “I liked actually getting to use software that professionals use! It was also nice to apply my own knowledge on something useful, it makes me remember what I’m learning more effortlessly and I enjoy it.” Thirty-eight students mentioned some aspect of the instruction as among their favorite features, including the topic lectures by the instructor or graduate students, or the survey activity itself.

**Table 1:**
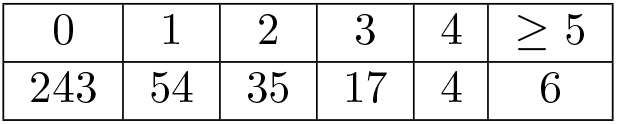
In how many classes at UCI (prior to this one) did you have the opportunity to apply the concepts you were learning about in class to a research project?

**Table 2:**
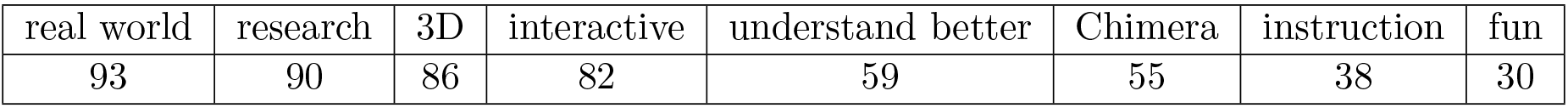
Please tell us what you liked best about the project. (topics from free response)

| real world | research | 3D | interactive | understand better | Chimera | instruction | fun |
| --- | --- | --- | --- | --- | --- | --- | --- |
| 93 | 90 | 86 | 82 | 59 | 55 | 38 | 30 |

Table 3 summarizes the most common responses given for what students liked least about the project. The most common responses focused on some aspect of the instructions being confusing or hard to follow (120 responses), or difficulty or frustration with the Chimera software (64 responses), although many also said that they got used to the software with practice. “What I liked least about the project was that the instructions were not always clear. While doing the survey during lecture time, I found myself confused by the instructions and I feel that affected the responses I submitted into the survey.” “I did not like having to download Chimera and go through that entire process for only a one time use.” “Getting used to using Chimera was my least favorite part, but it was also part of the learning experience.” “It was somewhat tough to get acquainted with the program in the beginning, but practice over the week helped with this.” Some students thought that the activity was rushed and they would have preferred either more class time or more time to work with their groups (11 responses). A few students did not like the open-ended nature of the assignment given that it is part of a live research project. Some were concerned about possibly providing incorrect information for the project (5 responses) or frustrated about not finding out the correct answer at the end (6 responses). “I didn’t like how stressful it was to think about how it could affect real research if we got a part incorrect.” “The right answer is not known.” “I wish I could have been able to look at other proteins gathered from the project to see what they looked like.” However, the total number of negative responses to participation in an active research project with no known answer (11 responses) were far outweighed by the positive ones described above (93 ‘real world experience’ + 90 ‘research,’ 183 total). Fifty-seven respondents specifically stated that they did not have a least favorite part, or that they liked everything about the exercise (blank responses were not included in this category). The least-liked aspects of the project included finding the instructions for using Chimera and comparing the two proteins confusing and feeling rushed to complete the activity during the class time. The most commonly given suggestions for improvement focused on making the instructions more clear. Another idea that was mentioned frequently was to allow students to analyze their proteins as a team. Finally, one student commented that the activity was difficult due to colorblindness, which is a useful reminder that instructions for changing the default colors in Chimera should be specifically discussed in the future.

**Table 3:**
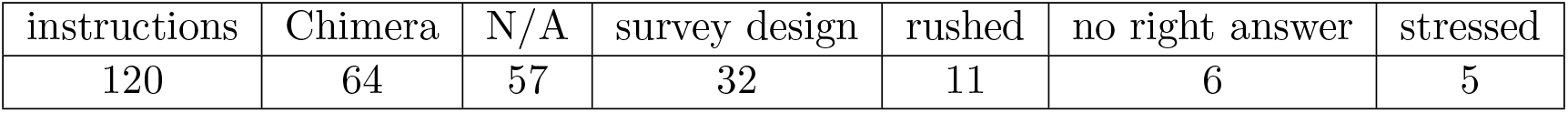
Please tell us what you liked least about the project. (topics from free response)

**Table 4:** Do you agree or disagree with the following statement: This research project helped me understand protein structure/function better. (choose one)

| strongly agree | agree | mildly agree | mildly disagree | disagree | strongly disagree |
| --- | --- | --- | --- | --- | --- |
| 72 | 170 | 96 | 11 | 5 | 5 |

**Table 5:** Do you agree or disagree with the following statement: This research project should continue to be a part of this course. (choose one)

| strongly agree | agree | mildly agree | mildly disagree | disagree | strongly disagree |
| --- | --- | --- | --- | --- | --- |
| 95 | 158 | 71 | 15 | 7 | 13 |

**Table 6:** How can this research project be improved? (topics from free response)

| clearer instructions | more feedback | Chimera video or demo |
| --- | --- | --- |
| 116 | 28 | 27 |

## CONCLUSION

This interactive exercise is adaptable for use in both smaller, upper-division and larger introductory biochemistry courses and can serve as an early exposure to current research projects; it could also be repeated after additional training with more advanced material. It enables students to use fundamental knowledge of protein secondary structures and motifs gained from lectures to actively build new skills that are essential for more advanced study and participation in research on structural biology and protein function. Student feedback following participation in the in-class activity was generally positive. In particular, students indicated that the potential for the work conducted in class to impact real-world research benefited their short-term engagement with the material and bolstered their sense of the value of investing in learning the information long-term. Criticism was primarily centered on actionable areas of improvement, such as providing more detailed instructions for using the software tools. We expect that future iterations will further benefit from tempering student expectations about the process and continuing to improve clarity in both the presentations and survey by conducting a separate analysis of how interpretations could lead to inconsistent answers. Increased participation and further development in this type of pedagogical tool will serve to not only improve students’ educational experience, but also expedite the pipeline for discovering new enzymes that are worthy of experimental validation, a particularly relevant activity in light of recent developments in protein structure prediction.

## Supporting information

Survey text and protein list

pdb files for example proteins

## ACKNOWLEDGMENTS

This research was supported by NSF award DMR-2002837 to R.W.M. and D.J. Tobias. M.A.S.-P. was supported by the HHMI Gilliam Fellowship for Advanced Study. M.G.C was supported by NIH award R25 GM055246 MBRS to the IMSD program at the University of California, Irvine. J.I.K. and B. N.-B. acknowledge support from the NSF GRFP. This material is based upon work supported by the National Science Foundation Graduate Research Fellowship under Grant No. DGE-1321846. Any opinion, findings, and conclusions or recommendations expressed in this material are those of the authors(s) and do not necessarily reflect the views of the National Science Foundation. The funders had no role in the design and conduct of the study, in the collection, analysis, and interpretation of the data, and in the preparation, review, or approval of the manuscript. We thank Carter Butts for advice about survey design. Most importantly, we gratefully acknowledge the hard work and helpful input of the students in Bio 98 (UC Irvine) and Chem341L (Fisk University).

## SUPPLEMENTAL MATERIAL

The SI contains the full text of the cysteine protease survey activity, the assessment survey, and a zip file containing the pdb files used in this exercise.

## AUTHOR CONTRIBUTIONS

R.W.M. and P.K. designed the course module concept. G.R.T., J.I.K., M.G.C., M.A.S.-P., J.L.U., S.M.D., R.W.M., and P.K. designed and taught the protein structure lecture material and activity training. J.I.K. 3D-printed structural models. S.M.D. and P.K. performed the course design and taught the classes. S.H.K., M.A.S.-P., J.I.K., G.R.T, M.G.C., B.N.-B., and R.W.M. designed the cysteine protease survey activity. J.I.K., P.K. and R.W.M. designed the assessment survey. R.W.M. analyzed the assessment survey data. F.S., V.F., G.R.T, J.I.K., and J.L.U. analyzed the cysteine protease survey data and provided the expert answers for each protein. J.L.U., F.S., V.F., G.R.T, M.G.C., E.M.D., and J.I.K. provided protein models and structure visualizations for the course materials and the manuscript. R.W.M., G.R.T., J.I.K. and P.K wrote the manuscript.

