## Supplementary material for "Course-based Undergraduate Research Module for Enzyme Discovery Using Protein Structure Prediction": Survey text and protein list

Cysteine Protease Survey Activity

### Welcome

Thank you for contributing to research on cysteine proteases! Your input will help us identify important functional features in newly-discovered digestive enzymes.

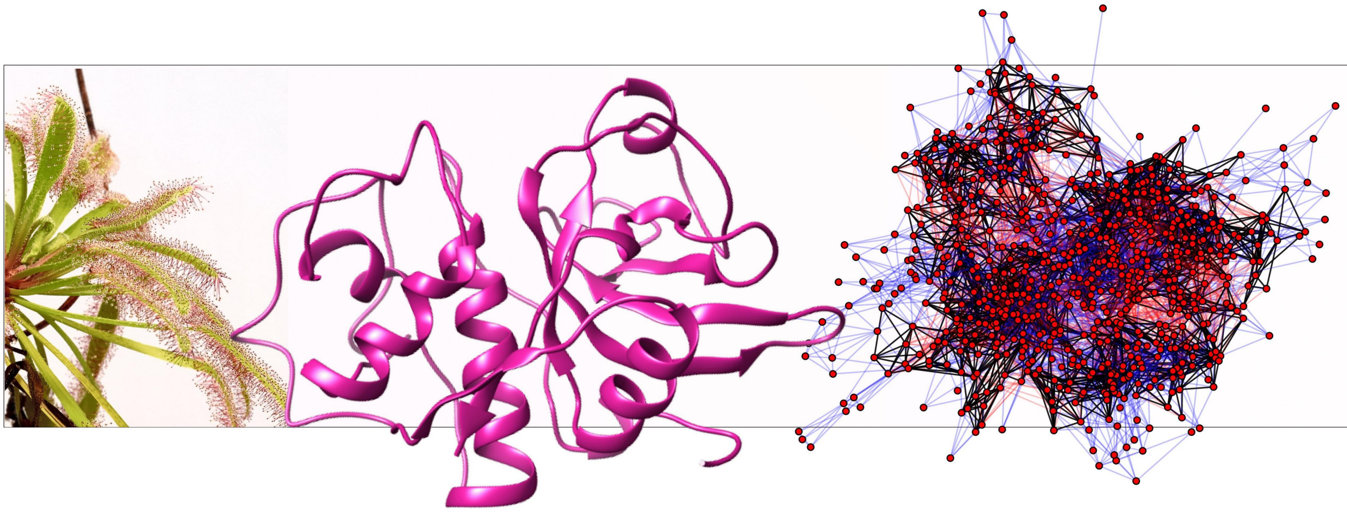

#### Setup

Download UCSF Chimera by going to https://www.cgl.ucsf.edu/chimera/download.html
Select the appropriate file for your computer's operating system and try to download the newest version possible

UCSF CHIMERA
an Extensible Molecular Modeling System

### Download Chimera

- Daily Builds
- Snapshot Releases
- Unsupported Releases
- Old Releases
- Bug Tracking System
- Licensing Information
- Experimental Chimera Features
- Plug-ins on the Web
- Graphics Driver Bugs
- Benchmark Results
- Chimera Source Code
- Cygwin Source Code

#### Current Production Releases

- See the release notes for a list of new features and other information.
- For more recent changes, use the snapshot and daily builds; they are less tested but usually reliable.
- **64-bit Releases:**

| Platform | Installer, Size, and Checksum | Date | Notes |
|---|---|---|---|
| Microsoft Windows 64-bit | chimera-1.12-win64.exe<br>Size: 150984696 bytes<br>MD5: b662f97cdacb8be458662aa638dd6694 | Oct 24, 2017 | Instructions<br>Documentation<br>Runs on Windows 7 or later. |
| Mac OS X 64-bit | chimera-1.12-mac64.dmg<br>Size: 131622631 bytes<br>MD5: 3a691632a3eba8b17286d23432bb6bbb | Oct 24, 2017 | Instructions<br>Documentation<br>Runs on Mac OS X 10.8 or later. |
| Linux 64-bit | chimera-1.12-linux_x86_64.bin<br>Size: 157795228 bytes<br>MD5: 24b9c2560e119988a3150b47c495737b | Oct 24, 2017 | Instructions<br>Documentation<br>Compiled on CentOS 5.11. |

- **32-bit Releases (for small memory computers):**

| Platform | Installer, Size, and Checksum | Date | Notes |
|---|---|---|---|
| Microsoft Windows | chimera-1.11.2-win32.exe<br>Size: 109260484 bytes<br>MD5: c0201380b0f328f57b9d839efcf646bd | Dec 02, 2016 | Instructions<br>Documentation<br>Runs on Windows 7 and 8 or later. |
| Mac OS X | chimera-1.11.2-mac.dmg<br>Size: 103774888 bytes<br>MD5: fa2ccd9c17c456d71088e81129c862d6 | Dec 02, 2016 | Instructions<br>Documentation<br>Runs on Mac OS X 10.8 or later. |
| Linux | chimera-1.11.2-linux.bin<br>Size: 119742278 bytes<br>MD5: 47dd12fbcbcfe01ea678599dd7001a6b | Dec 02, 2016 | Instructions<br>Documentation<br>Compiled on Debian 4 (etch). |

Enter the number of the protein that you were assigned below.

**Alignment**

1. Open UCSF Chimera
2. Open the .pdb file for the reference sequence to generate a model of the protein. For cysteine proteases, this will be papain(PAPA1_CARPA). You should get an image like the one below. The reference sequence will be colored beige

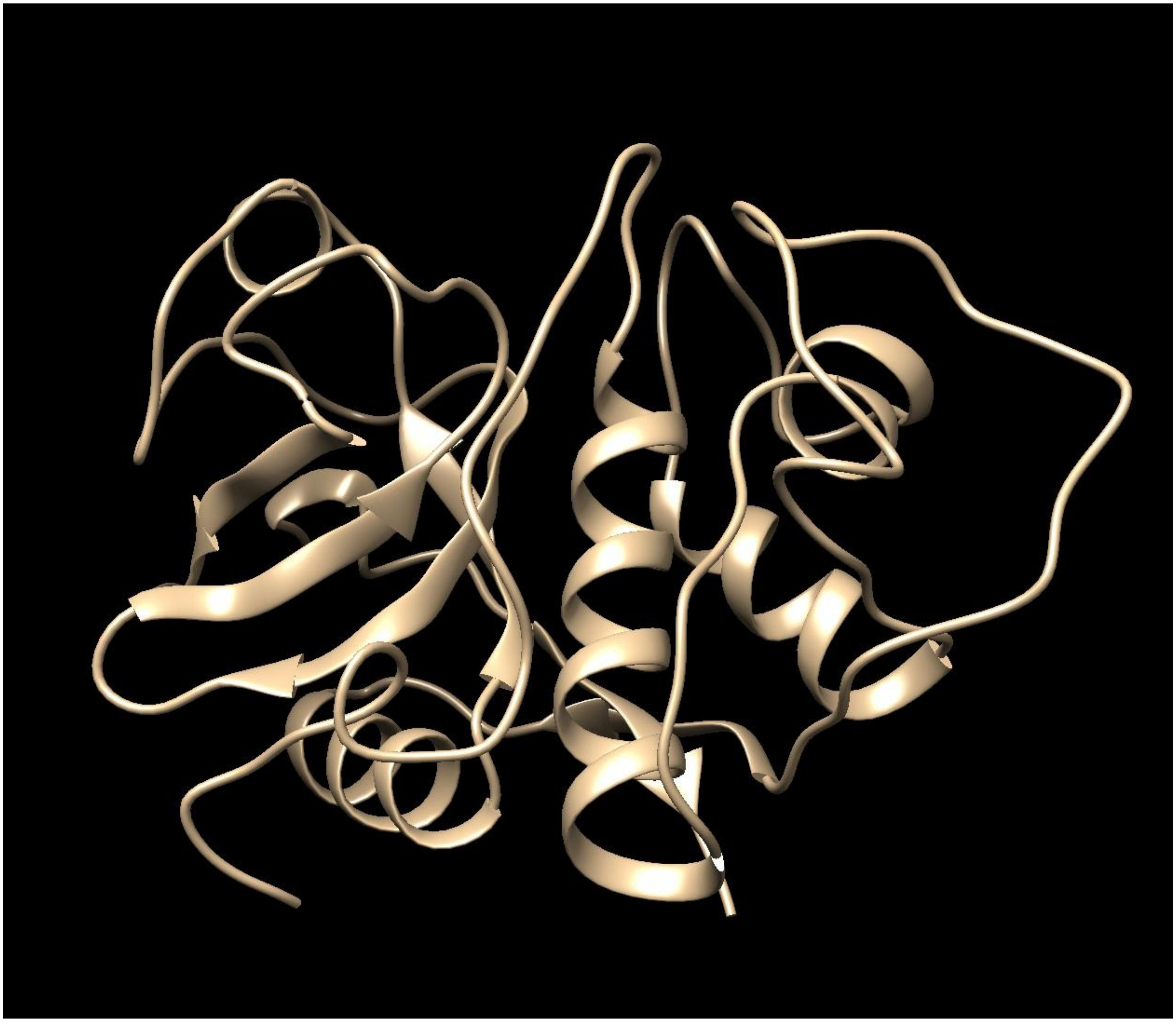

3. Open a second protein, the one you will be comparing to papain. You will now have two proteins on top of one another in no particular orientation. Your protein will be colored light blue.
4. In order to align the two proteins with one another, scroll to tools. Then select the following Tools –– Structure Comparison –– Matchmaker. A screen like the one below will be generated. Select PAPA1_CARPA_cut.eq.1.pdb (#0) for your reference structure, and the protein of comparison as the structure to match. Keep all the other defaults the same, and then hit "Apply.

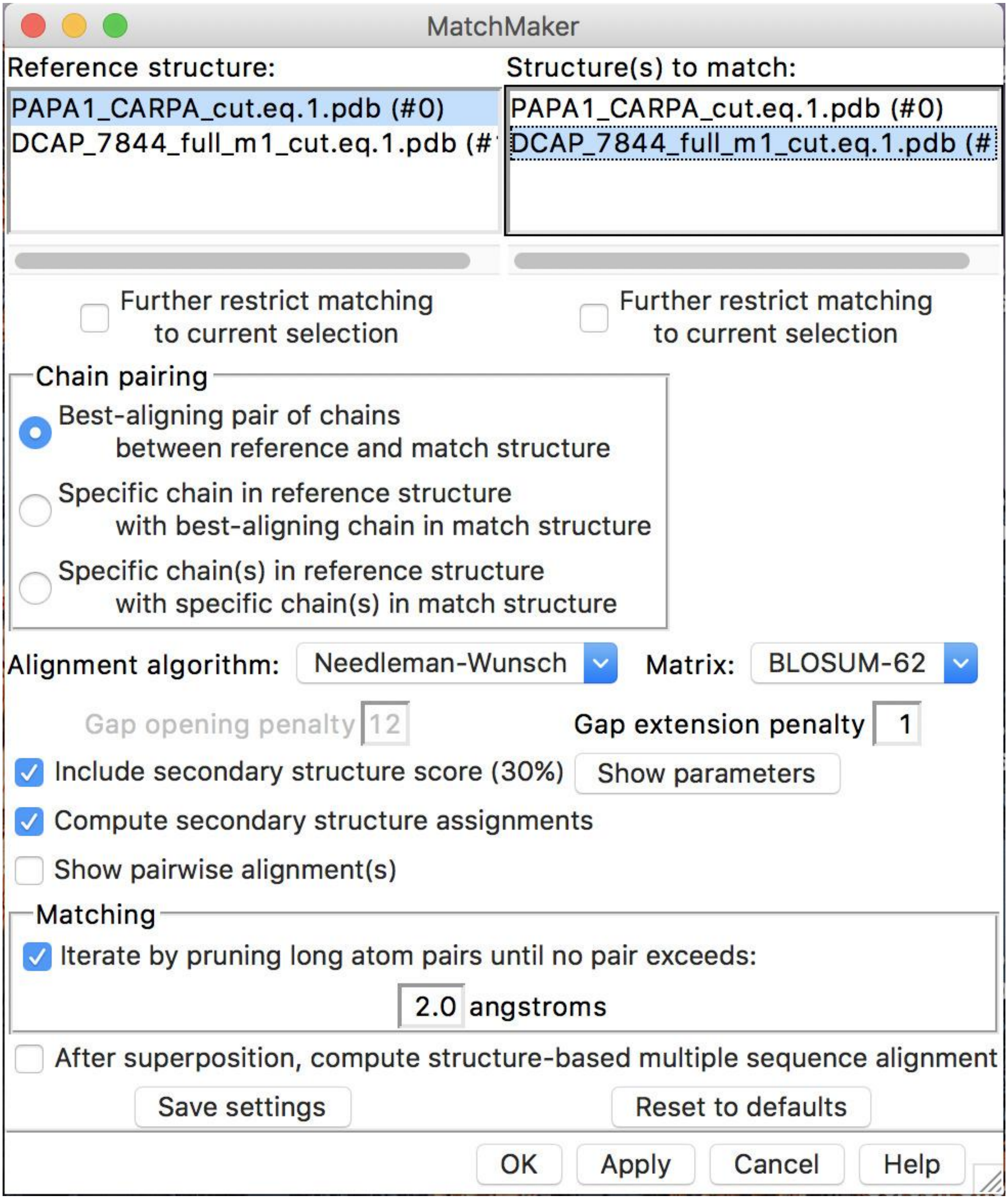
MatchMaker
Reference structure:
Structure(s) to match:
PAPA1_CARPA_cut.eq.1.pdb (#0)
DCAP_7844_full_m1_cut.eq.1.pdb (#
PAPA1_CARPA_cut.eq.1.pdb (#0)
DCAP_7844_full_m1_cut.eq.1.pdb (#
Further restrict matching to current selection
Further restrict matching to current selection
Chain pairing
Best-aligning pair of chains between reference and match structure
Specific chain in reference structure with best-aligning chain in match structure
Specific chain(s) in reference structure with specific chain(s) in match structure
Alignment algorithm: Needleman-Wunsch
Matrix: BLOSUM-62
Gap opening penalty 12
Gap extension penalty 1
Include secondary structure score (30%)
Show parameters
Compute secondary structure assignments
Show pairwise alignment(s)
Matching
Iterate by pruning long atom pairs until no pair exceeds:
2.0 angstroms
After superposition, compute structure-based multiple sequence alignment
Save settings
Reset to defaults
OK
Apply
Cancel
Help

Your protein sequences will now be aligned!
5. Open the amino acid sequences for the proteins. Go to Tools –– Sequence –– Sequence and you will reach the screen below. Open both protein sequences.

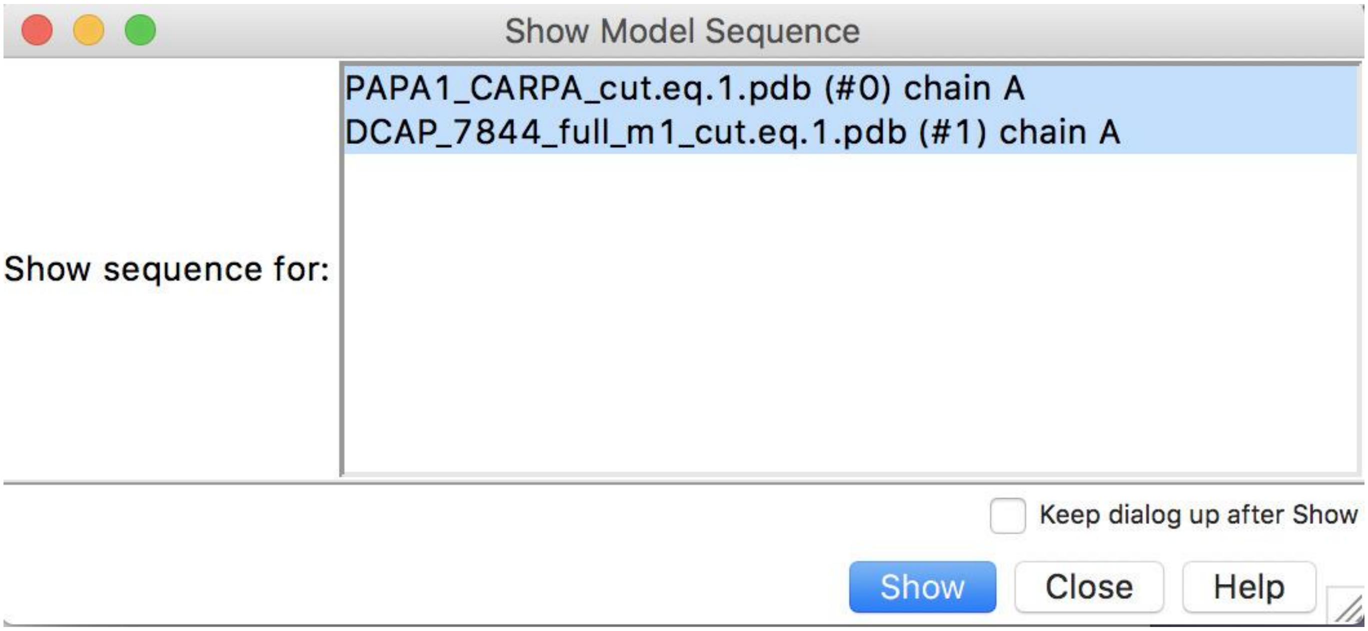

You should get a pop-up that looks like this. The yellow highlights mean that there is an alpha helix in that area and the green highlights
correspond to beta strands.

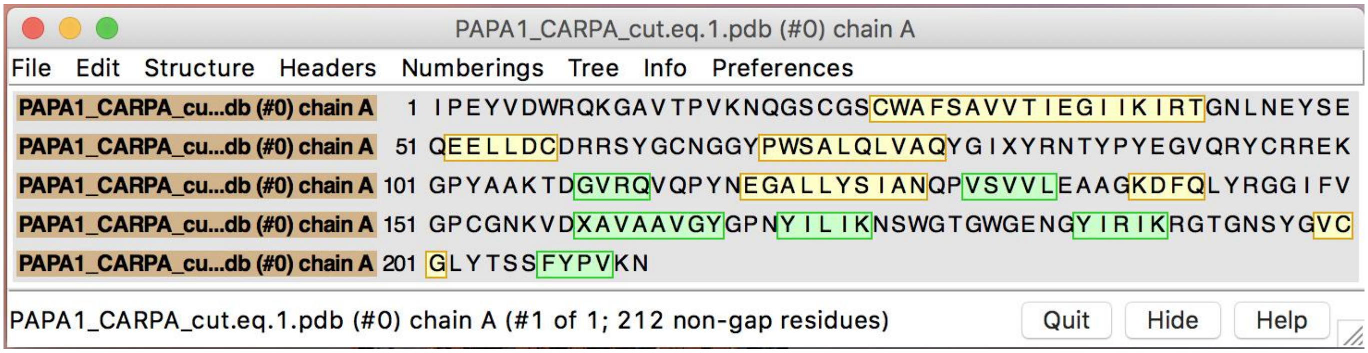

### General Questions

#### General Questions

For each question below you will compare your assigned structure to the structure of papain

How many amino acids long is your protein? (Hover the cursor over the last amino acid in the sequence and the menu will display the residue number)

| | 100 | 130 | 160 | 190 | 220 | 250 | 280 | 310 | 340 | 370 | 400 |
|---|---|---|---|---|---|---|---|---|---|---|---|
| Total number of amino acids | | | | | | | | | | | |

The tertiary structure of the protein below contains secondary structure elements. The green colored part of the structure is a beta strand, while the yellow colored part is an alpha helix. You can highlight an alpha helix in your protein by selecting the corresponding residues in your protein. Then go to Actions –– Color and then select the color you want to highlight that part of your structure.

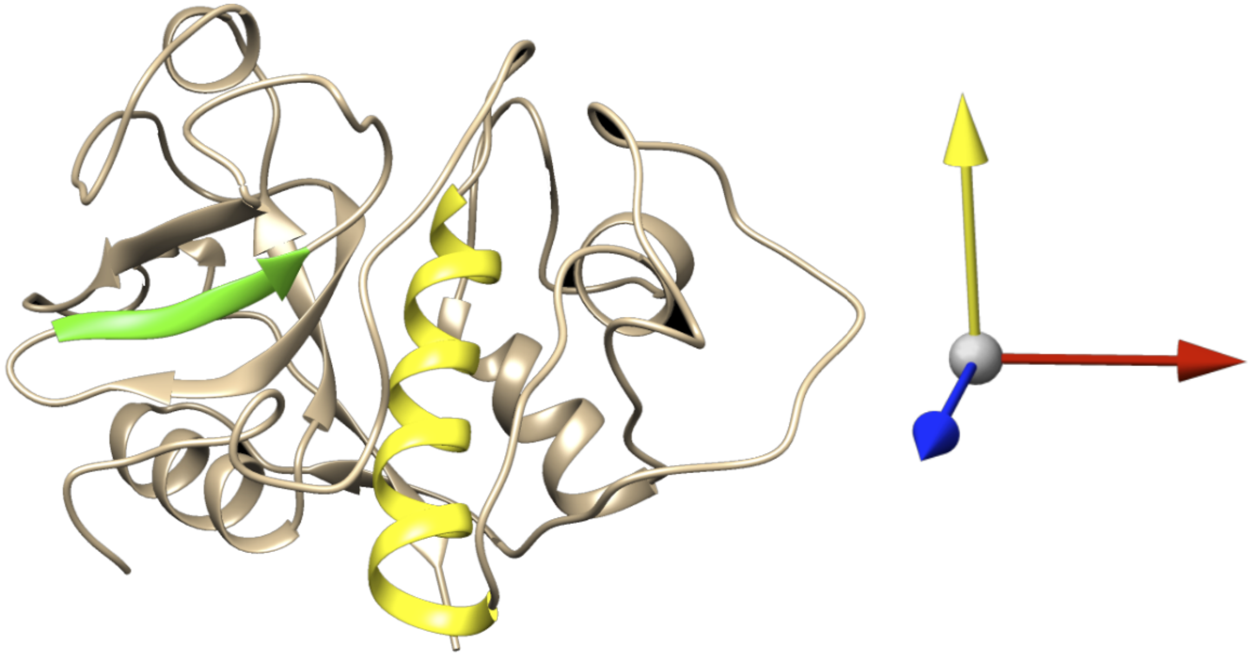

#### Alpha Helix Analysis

---

We are going to compare the first alpha helix in papain to the corresponding helix in your protein. First select the first alpha helix in papain. The selected area will be highlighted green in your Chimera session for easy viewing. In the example below, the first alpha helix has been colored yellow. Color this helix in papain.

Highlight the corresponding alpha helix in your protein, make it a different color, and take a screenshot like in the example below.

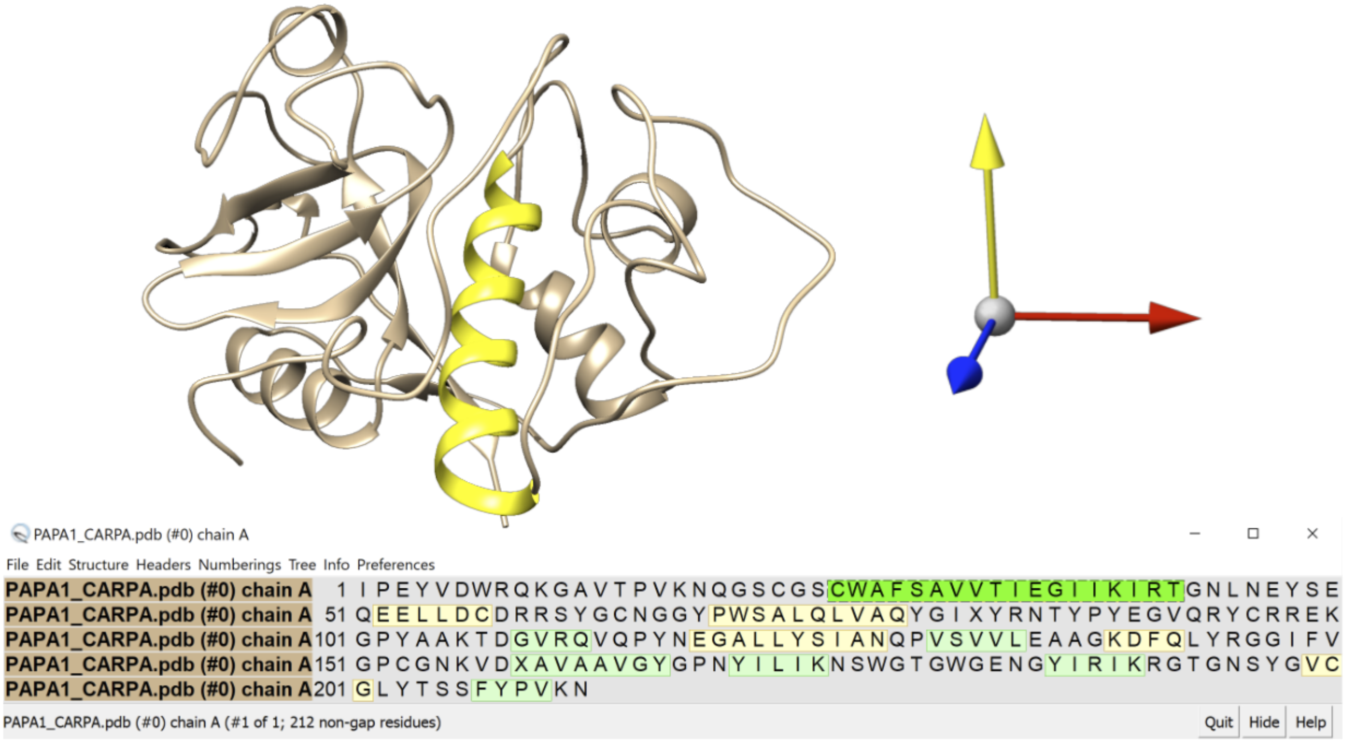

In the example below papain (beige) is overlaid on a protein similar to one that is assigned to you (blue). Here the first alpha helix on the blue structure has been highlighted and colored orange.

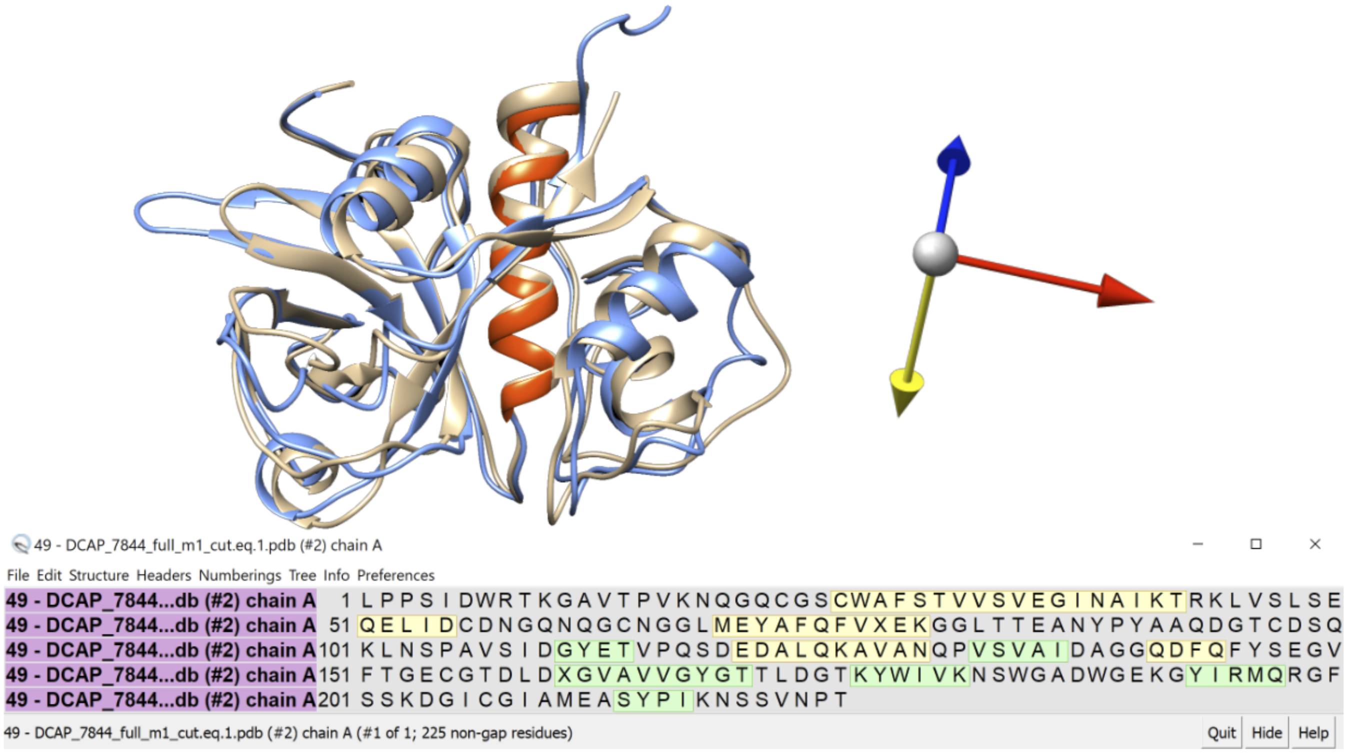

Remember the first alpha helix will be the first section highlighted in yellow from the left in the sequence of papain. You can use the mouse to highlight that section of the sequence to select it and change the color of those amino acids on the structure. In your structure, there might be several helices that appear in the amino acid sequence before one that aligns in space with the first helix in papain in the 3D structures. Ignore those helices and find a helix in your structure that lines up with the first helix in papain in space. Is there a helix on your structure that lines up with the first helix in papain?

Yes

No

For each of the remaining alpha helices in **papain:**

Determine if your protein has a corresponding alpha helix in the structure. Make sure to compare the location of the helices in the structure.

If your structure has more alpha helices than papain, disregard them for this question, but note this for the last question. But make sure to only look at the ones that line up with the papain structure. For example, the second helix in papain might align with the fifth helix in your structure. If you see something like that then you would answer "Yes" to "alpha helix 2" below. If no helix in your structure aligns with the second helix in papain then you would answer "No".

| | Yes | No |
|---|---|---|
| alpha helix 2 | ○ | ○ |
| alpha helix 3 | ○ | ○ |
| alpha helix 4 | ○ | ○ |
| alpha helix 5 | ○ | ○ |

| | Yes | No |
| --- | --- | --- |
| alpha helix 6 | ○ | ○ |

### Block 2

#### Alpha Helix Analysis Continued

In Chimera there are two methods for determining residue numbers when looking at the protein structure. You can open the sequence menu and count to where the residue is. Or, in the second, simpler method, you can hover your mouse over a certain part of the protein and the number will pop up, giving the type of amino acid, followed by the residue number. In the example below we have selected Threonine 33. This THR is near the middle of the first alpha helix and is residue number 33 in the reference structure papain.

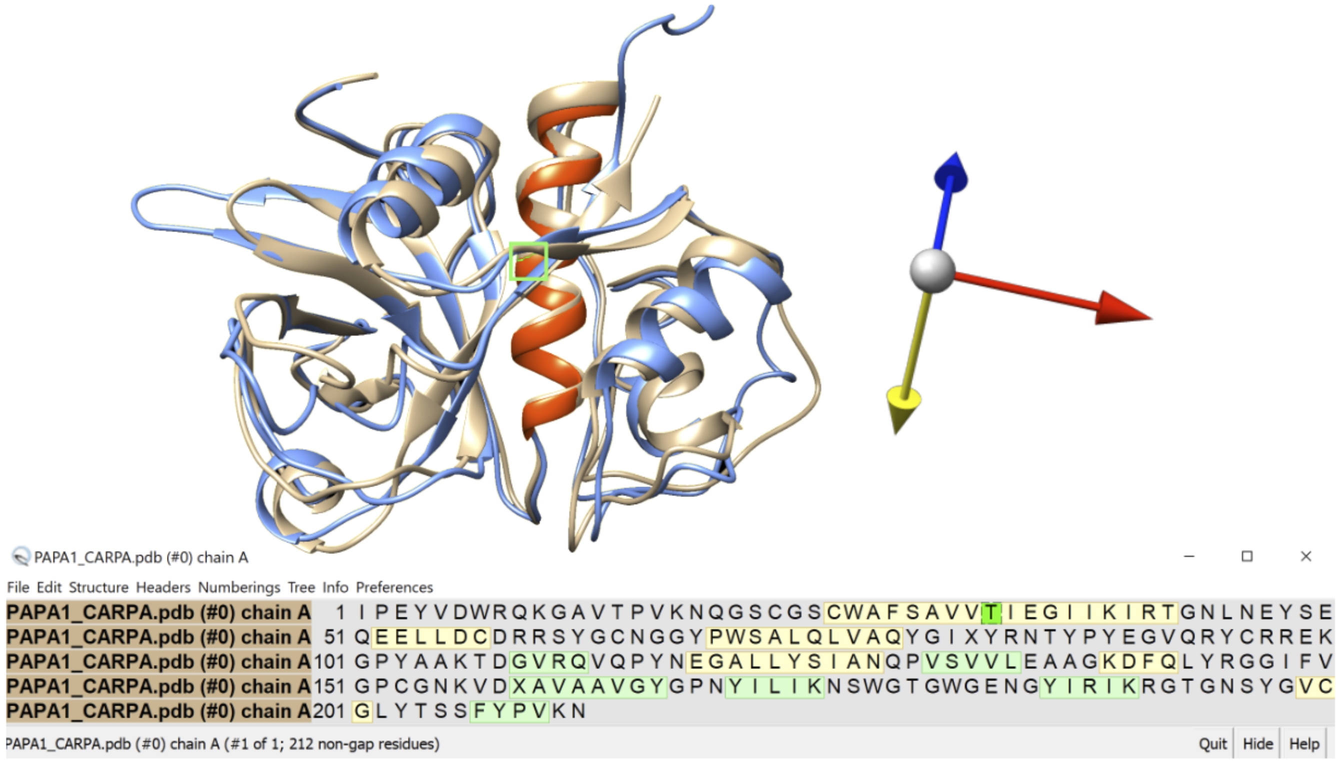

Select a residue in the alpha helix number 1 of your assigned protein (the blue one) that is close to the center. In the example shown above for the reference sequence papain, this is amino acid number 33, the green highlighted threonine. (The number will probably not be the same as in papain.)

1 11 21 31 41 51 60 70 80 90 100

Center reside of alpha helix 1

Now do the same for the rest of the alpha helices. Select a residue that is in the middle of the helix and enter the number for that residue. Remember if your protein has more helices than papain ignore the extra ones for now, but note this for the last question. Below you should only see alpha helices for which you answered "Yes" above.

| | 1 | 41 | 81 | 121 | 161 | 201 | 240 | 280 | 320 | 360 | 400 | |
|---|---|---|---|---|---|---|---|---|---|---|---|---|
| alpha helix 2 | | | | | | | | | | | | ☐ |
| alpha helix 3 | | | | | | | | | | | | ☐ |
| alpha helix 4 | | | | | | | | | | | | ☐ |
| alpha helix 5 | | | | | | | | | | | | ☐ |
| alpha helix 6 | | | | | | | | | | | | ☐ |

#### Block 3

##### Beta Strands

The following image illustrates the position of the first beta strand in papain, it is colored in green.

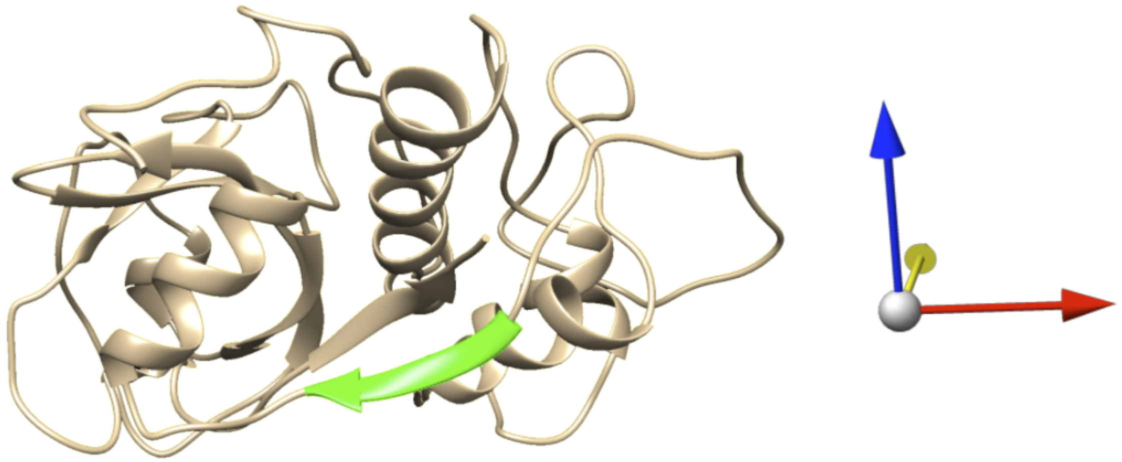

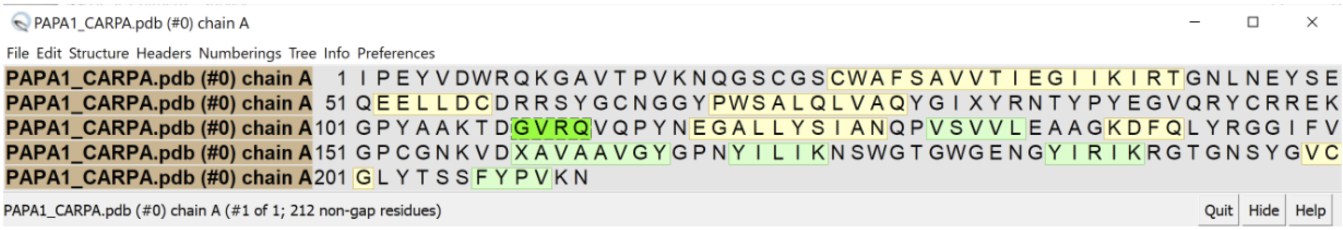

In the example below papain (beige) is overlaid on a protein similar to one assigned to you (blue). Highlighted in green is the first beta strand on the blue structure that corresponds to one in papain.

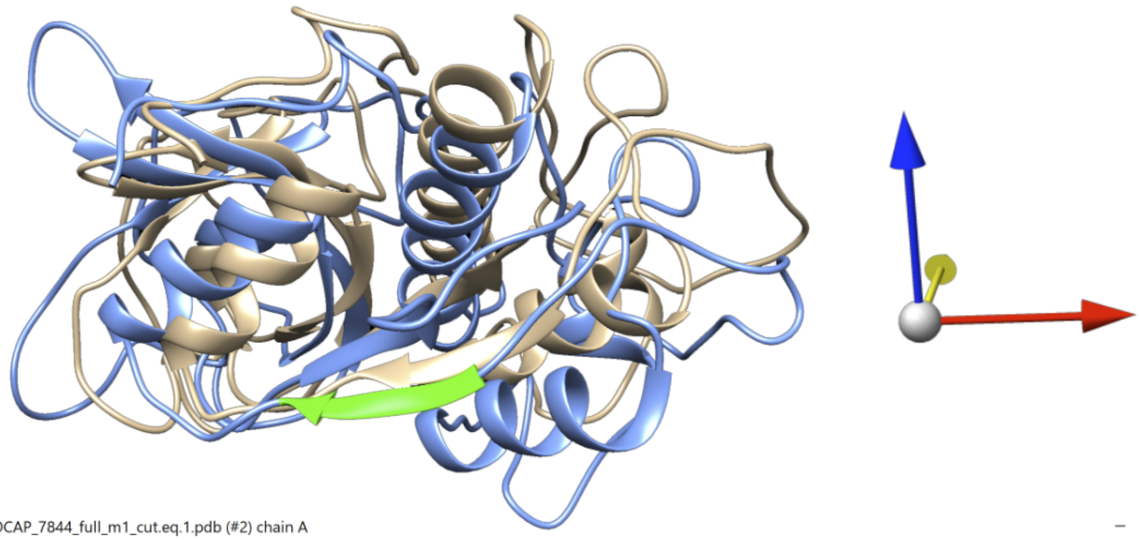

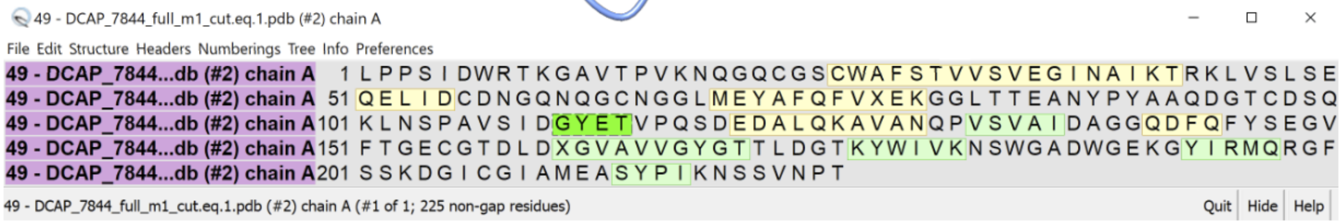

Remember the first beta strand will be the first section highlighted in green from the left in the sequence of papain. You can use the mouse to highlight that section of the sequence to select it and change the color of those amino acids on the structure.

Now highlight the first beta strand in your assigned protein.

Does it line up with the first beta strand in papain.

Yes

No

Now do the same for the rest of the beta strands in papain.

| | Yes | No |
|---|---|---|
| beta strand 2 | ○ | ○ |
| beta strand 3 | ○ | ○ |
| beta strand 4 | ○ | ○ |
| beta strand 5 | ○ | ○ |
| beta strand 6 | ○ | ○ |

#### Block 4

##### Beta Strand Analysis Continued

---

Below is an example that shows a selected residue near the middle of the first beta strand. The residue selected is Valine and the residue number is 110.

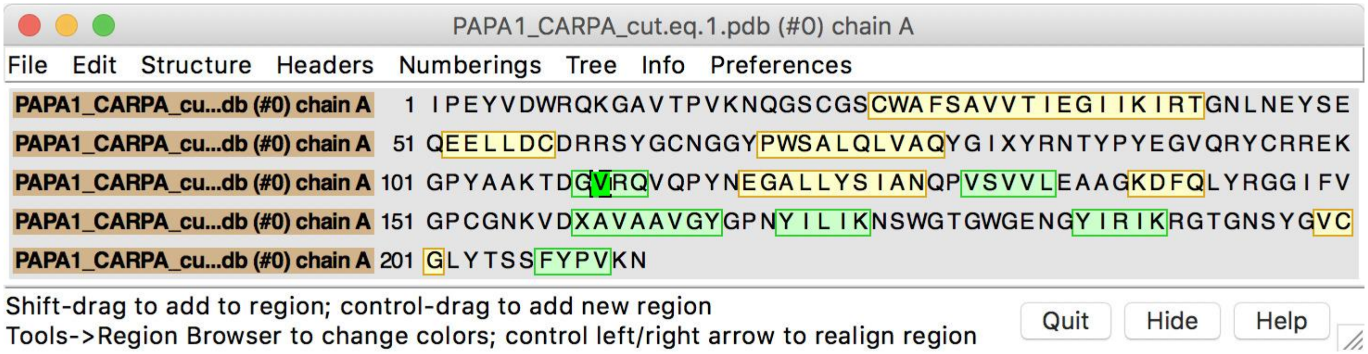

Select a residue in the beta strand number 1 that is close to the center (like residue number 110 in the example above). What number (##) is that residue in your protein's sequence?

| | 1 | 21 | 41 | 61 | 81 | 101 | 120 | 140 | 160 | 180 | 200 |
|---|---|---|---|---|---|---|---|---|---|---|---|
| Center of beta strand 1 | | | | | | | | | | | |

Do the same for the rest of the beta strands. Below you should only see beta strands for which you answered "Yes" above.

| | 1 | 41 | 81 | 121 | 161 | 201 | 240 | 280 | 320 | 360 | 400 |
|---|---|---|---|---|---|---|---|---|---|---|---|
| Beta strand 2 | | | | | | | | | | | |
| beta strand 3 | | | | | | | | | | | |
| beta strand 4 | | | | | | | | | | | |

| | 1 | 41 | 81 | 121 | 161 | 201 | 240 | 280 | 320 | 360 | 400 |
|---|---|---|---|---|---|---|---|---|---|---|---|
| beta strand 5 | | | | | | | | | | | |
| beta strand 6 | | | | | | | | | | | |

#### Block 5

##### Beta Sheet analysis

A beta sheet is made up of beta strands like the area of papain highlighted in light green below.

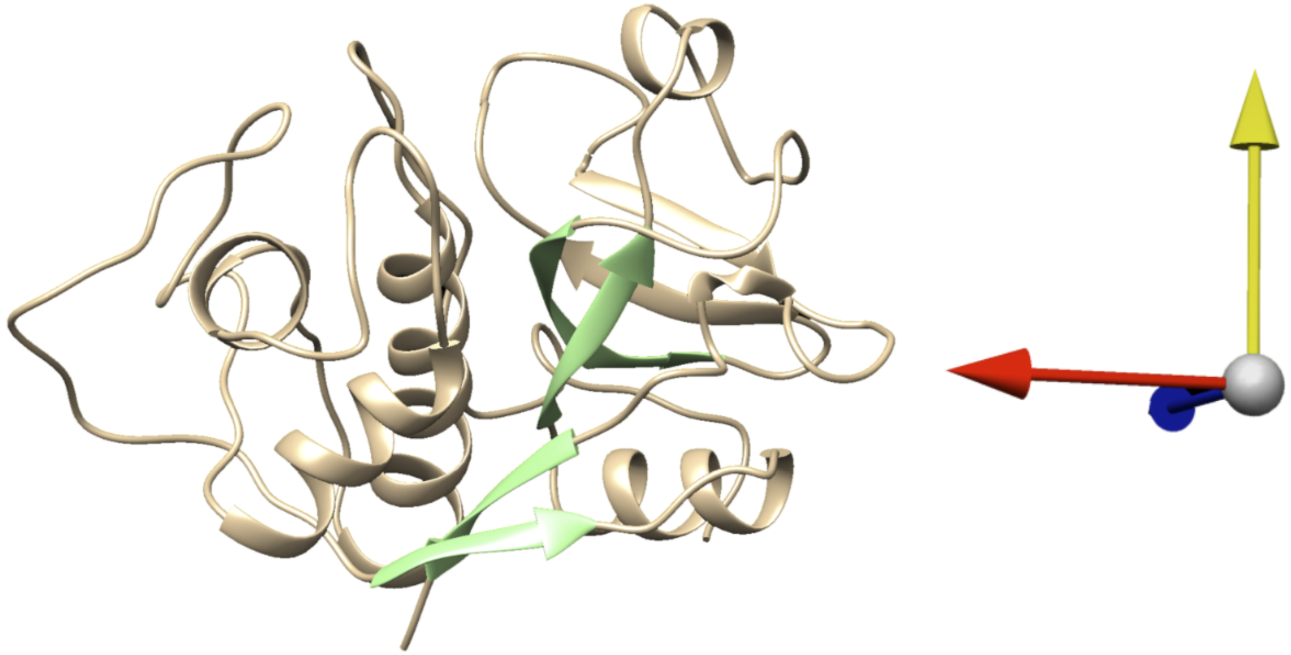

Below is an example of this beta sheet area selected in magenta on a protein like the one assigned to you. You can select multiple strands at the same time by holding shift as you drag your cursor across the sequence.

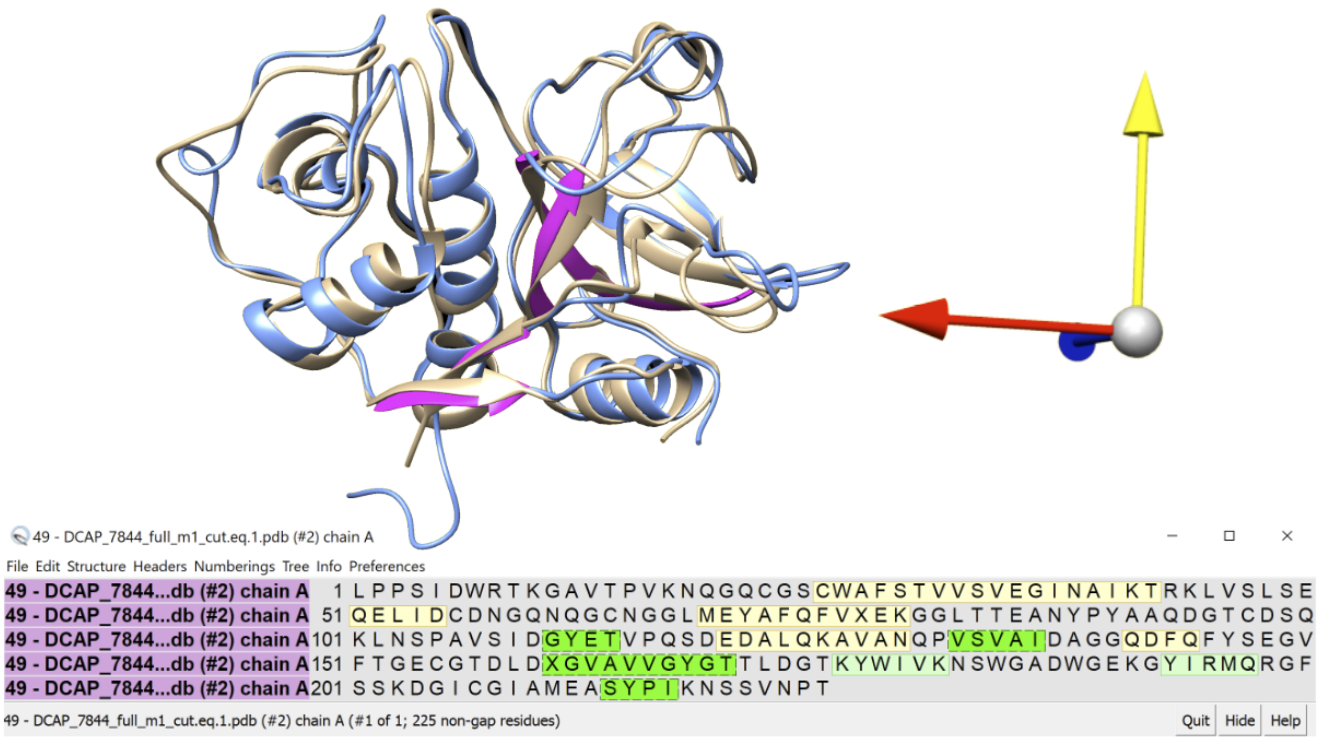

#### Does your protein have a beta sheet like the one seen in papain?

Yes, the beta sheet is in my protein and it is made up of four beta strands

Yes, the beta sheet is in my protein but is not made of four beta strands

No, this beta sheet is not in my protein

#### The two beta strands colored in dark green below make up a different beta sheet that is also a key part of the secondary structure of papain.

---

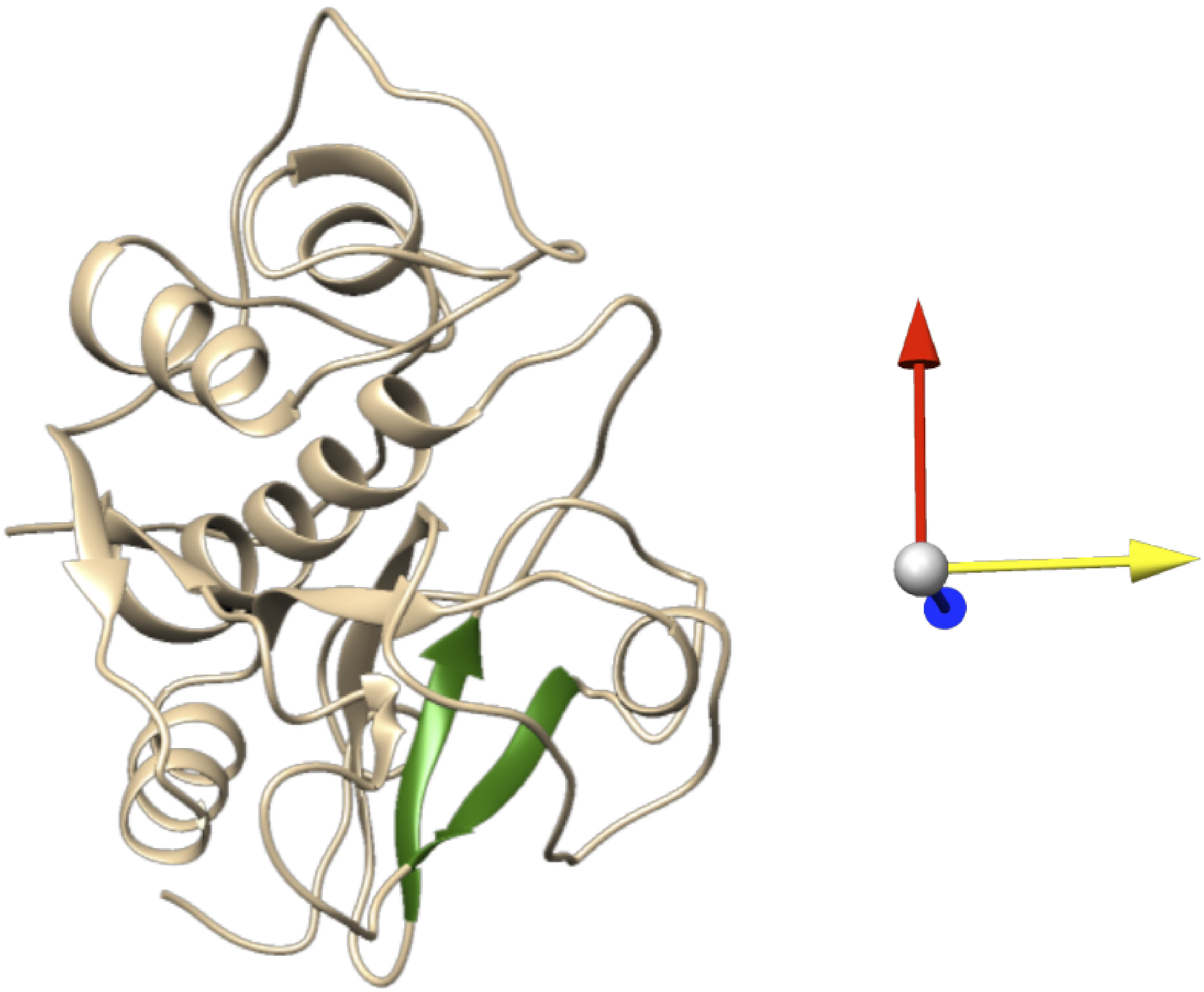

Below is an example of this beta sheet area selected in red on a protein like the one assigned to you.

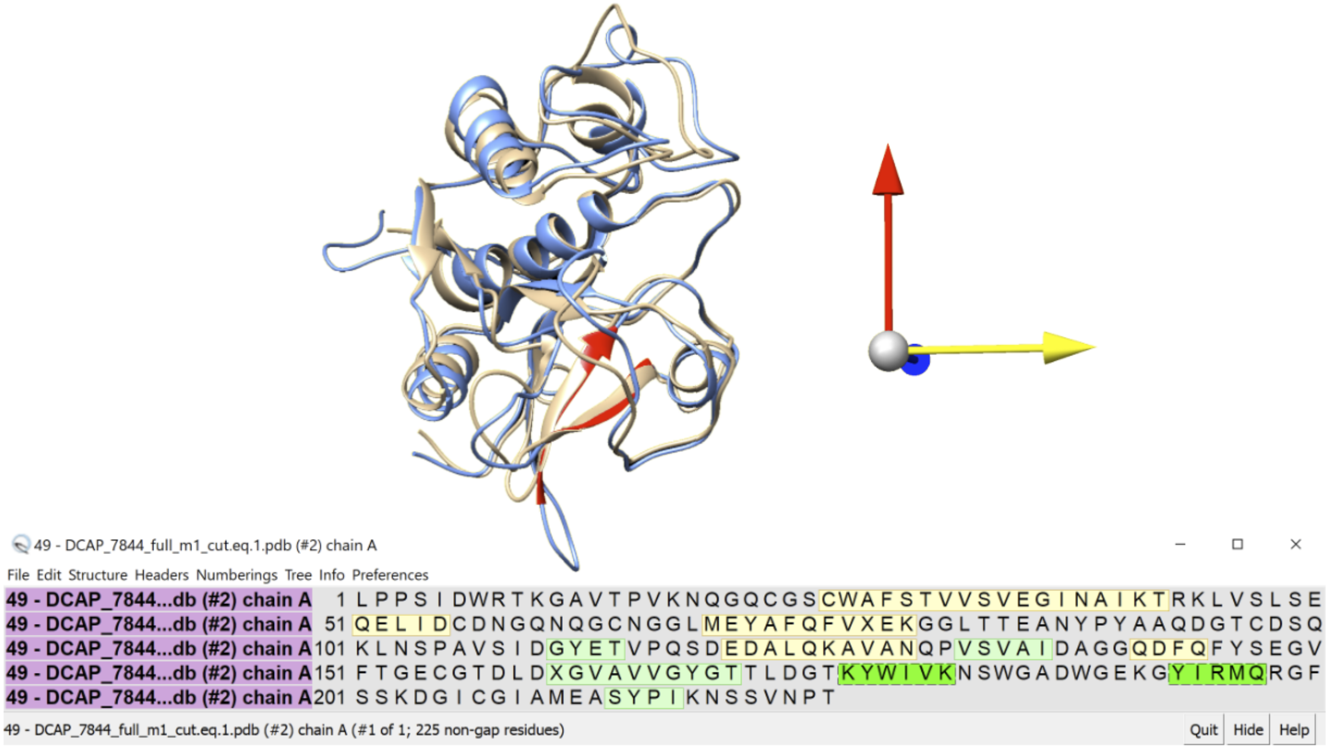

#### Does your protein have a beta sheet like the one seen in papain? (Note whether or not there are the same number of beta strands in the beta sheet)

Yes, the beta sheet is in my protein and it is made up of two beta strands

Yes, the beta sheet is in my protein but is not made of two beta strands

No, This beta sheet is not is my protein

### Block 6

#### Active Site Analysis

The active site of papain is the catalytic triad of Cys-25, Hsp-159, and Asn-175. Below the atom representation of the sidechains on these residues are shown for clarity. To do this yourself, highlight residues Cys-25, Hsp-159, and Asn-175 in papain's sequence, then go to Actions –– Atoms/Bonds –– show. While the active site may not be exactly the same for your protein, this is a good starting point. The green arrow points to "V-Shaped" area known as the active site cleft of papain.

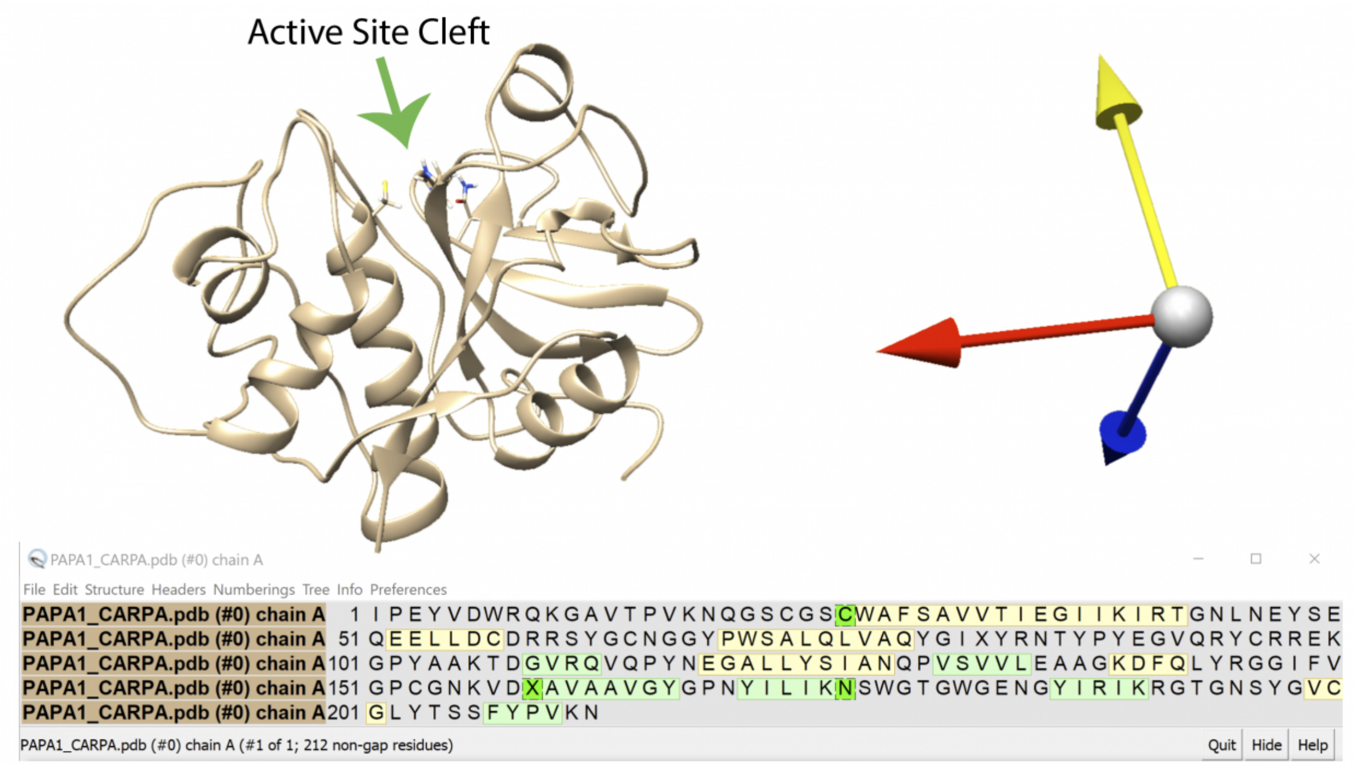

Some proteins have features that will fully or partially block the active site. Below is an example of such a feature. The purple protein (DCAP_2263) has a very long N-terminus and loop that is not in papain (beige). These features are shown in the magenta dashed circles.

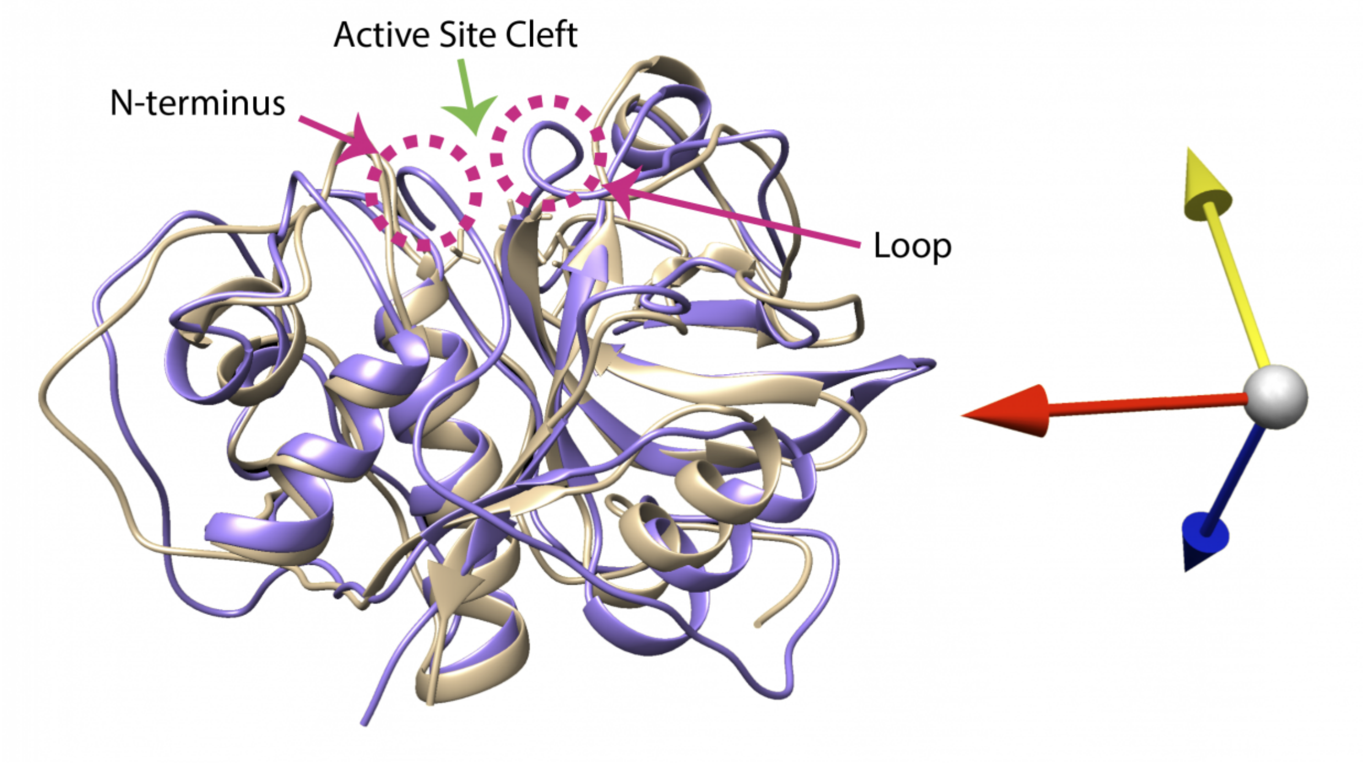

Do you see a feature that is partially or fully blocking the active site of your protein? (Many proteins do not have one)

Yes

No

If you do have a feature that blocks the active site in your protein then select a residue that is on that feature and enter the number of that residue below. If there is more than one feature such as above, choose the one closest to the cleft (in this case the loop) and make note of the other for the last question.

| | 1 | 41 | 81 | 121 | 161 | 201 | 240 | 280 | 320 | 360 | 400 |
|---|---|---|---|---|---|---|---|---|---|---|---|
| Click to write Choice 1 | | | | | | | | | | | |

#### Block 7

##### N-terminus and C-terminus

---

The following image show the N-terminus and C-terminus of the two proteins. In this example, the N-termini are the same length, while the C-terminus of the blue protein is longer.

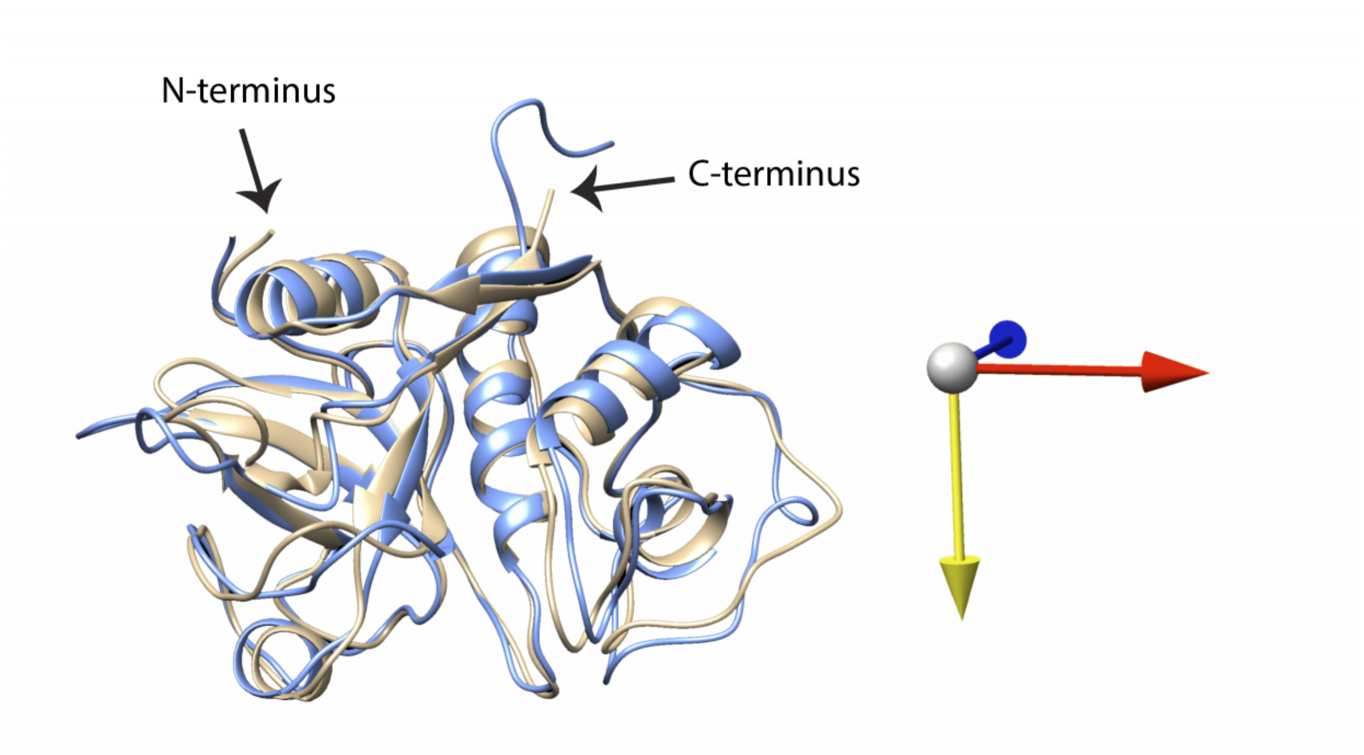

Is the N-terminus of your protein longer, shorter, or the same length as the N-terminus of papain?

Longer

Shorter

The same length

Is the C-terminus of your protein longer, shorter, or the same as the C-terminus of papain?

Longer

Shorter

The same length

### Block 8

#### Wrapping up

Previously in this survey you have answered questions on the overall amino acid length, secondary structure features such as the overlap of alpha helices and beta sheets with the reference structure, features around the active site cleft, and lengths of the termini for your assigned protein.

Review the structure for elements discussed in the lecture and from your previous studies. What differences does your protein have when compared to papain that were either not fully captured (such as more helices or strands) or not addressed at all in earlier questions? Describe the feature(s).

Do you have any suggestions to improve this survey?

***Please highlight the most interesting part of your protein, save that image as a JPEG, and upload that image to your Canvas assignment.

**This is how you get credit for the assignment!**

Thank you for completing the survey! You are contributing to cysteine protease research!

Powered by Qualtrics

Table S1: Available pdb files.

| Protein Name | UniProt[1] ID | Organism | Strain Info. | File Name | Citation | University_ Course |
|---|---|---|---|---|---|---|
| **E3DLQ8** | E3DLQ8_HALPG | *Halanaerobium praevalens* | ATCC 33744 / DSM 2228 / GSL | 1.pdb | 2 | UCI_Bio98 |
| **A0A1Q3BJ77** | A0A1Q3BJ77_CEPFO | *Cephalotus follicularis* | cv. St1 | 2.pdb | 3 | UCI_Bio98 |
| **A0A1Q3BIX5** | A0A1Q3BIX5_CEPFO | *Cephalotus follicularis* | cv. St1 | 3.pdb | 3 | UCI_Bio98 |
| **A0A1Q3BV84** | A0A1Q3BV84_CEPFO | *Cephalotus follicularis* | cv. St1 | 4.pdb | 3 | UCI_Bio98 |
| **A0A1Q3CR93** | A0A1Q3CR93_CEPFO | *Cephalotus follicularis* | cv. St1 | 5.pdb | 3 | UCI_Bio98 |
| **A0A1Q3CMQ8** | A0A1Q3CMQ8_CEPFO | *Cephalotus follicularis* | cv. St1 | 6.pdb | 3 | UCI_Bio98 |
| **A0A1Q3AVM1** | A0A1Q3AVM1_CEPFO | *Cephalotus follicularis* | cv. St1 | 7.pdb | 3 | UCI_Bio98 |
| **A0A1Q3BQA4** | A0A1Q3BQA4_CEPFO | *Cephalotus follicularis* | cv. St1 | 8.pdb | 3 | UCI_Bio98 |
| **A0A1Q3B3S8** | A0A1Q3B3S8_CEPFO | *Cephalotus follicularis* | cv. St1 | 9.pdb | 3 | UCI_Bio98 |
| **A0A1Q3DBM0** | A0A1Q3DBM0_CEPFO | *Cephalotus follicularis* | cv. St1 | 10.pdb | 3 | UCI_Bio98 |
| **A0A1Q3AYB2** | A0A1Q3AYB2_CEPFO | *Cephalotus follicularis* | cv. St1 | 11.pdb | 3 | UCI_Bio98 |
| **A0A1Q3BXS8** | A0A1Q3BXS8_CEPFO | *Cephalotus follicularis* | cv. St1 | 12.pdb | 3 | UCI_Bio98 |
| **A0A1Q3B305** | A0A1Q3B305_CEPFO | *Cephalotus follicularis* | cv. St1 | 13.pdb | 3 | UCI_Bio98 |
| **A0A1Q3BAZ0** | A0A1Q3BAZ0_CEPFO | *Cephalotus follicularis* | cv. St1 | 14.pdb | 3 | UCI_Bio98 |
| **A0A1Q3D7H5** | A0A1Q3D7H5_CEPFO | *Cephalotus follicularis* | cv. St1 | 15.pdb | 3 | UCI_Bio98 |
| **A0A1Q3C6Y1** | A0A1Q3C6Y1_CEPFO | *Cephalotus follicularis* | cv. St1 | 16.pdb | 3 | UCI_Bio98 |
| **A0A1Q3CNW3** | A0A1Q3CNW3_CEPFO | *Cephalotus follicularis* | cv. St1 | 17.pdb | 3 | UCI_Bio98 |
| **DCAP_0302** | NA | *Drosera capensis* | wide | 18.pdb | 4 | UCI_Bio98 |
| **DCAP_5945** | NA | *Drosera capensis* | wide | 19.pdb | 4 | UCI_Bio98 |
| **DCAP_6097** | NA | *Drosera capensis* | wide | 20.pdb | 4 | UCI_Bio98 |
| **DCAP_6547** | NA | *Drosera capensis* | wide | 21.pdb | 4 | UCI_Bio98 |
| **DCAP_7518** | NA | *Drosera capensis* | wide | 22.pdb | 4 | UCI_Bio98 |
| **DCAP_0624** | NA | *Drosera capensis* | wide | 23.pdb | 4 | UCI_Bio98 |
| **DCAP_0995** | NA | *Drosera capensis* | wide | 24.pdb | 4 | UCI_Bio98 |
| **DCAP_1658** | NA | *Drosera capensis* | wide | 25.pdb | 4 | UCI_Bio98 |
| **DCAP_2190** | NA | *Drosera capensis* | wide | 26.pdb | 4 | UCI_Bio98 |
| **DCAP_3192** | NA | *Drosera capensis* | wide | 27.pdb | 4 | UCI_Bio98 |
| **DCAP_3193** | NA | *Drosera capensis* | wide | 28.pdb | 4 | UCI_Bio98 |
| **DCAP_3872** | NA | *Drosera capensis* | wide | 29.pdb | 4 | UCI_Bio98 |
| **DCAP_4240** | NA | *Drosera capensis* | wide | 30.pdb | 4 | UCI_Bio98 |
| **DCAP_3968** | NA | *Drosera capensis* | wide | 31.pdb | 4 | UCI_Bio98 |

| | | | | | | |
|---|---|---|---|---|---|---|
| **DCAP_4793** | NA | *Drosera capensis* | wide | 32.pdb | 4 | UCI_Bio98 |
| **DCAP_4952** | NA | *Drosera capensis* | wide | 33.pdb | 4 | UCI_Bio98 |
| **DCAP_5115** | NA | *Drosera capensis* | wide | 34.pdb | 4 | UCI_Bio98 |
| **DCAP_2570** | NA | *Drosera capensis* | wide | 1 - DCAP_2570_full_m1_cut.eq.1.pdb | 4 | Fisk_Chem 341L |
| **DCAP_2555** | NA | *Drosera capensis* | wide | 2 - DCAP_2555_full_m1_cut.eq.1.pdb | 4 | Fisk_Chem 341L |
| **DCAP_5561** | NA | *Drosera capensis* | wide | 3 - DCAP_5561_full_m1_cut.eq.1.pdb | 4 | Fisk_Chem 341L |
| **DCAP_5667** | NA | *Drosera capensis* | wide | 4 - DCAP_5667_full_m1_cut.eq.1.pdb | 4 | Fisk_Chem 341L |
| **DCAP_6618** | NA | *Drosera capensis* | wide | 5 - DCAP_6618_full_m1_cut.eq.1.pdb | 4 | Fisk_Chem 341L |
| **DCAP_7518** | NA | *Drosera capensis* | wide | 6 - DCAP_7518_nogranulin_m1_cut.eq.1.pdb | 4 | Fisk_Chem 341L |
| **DCAP_7656** | NA | *Drosera capensis* | wide | 7 - DCAP_7656_full_m1_cut.eq.1.pdb | 4 | Fisk_Chem 341L |
| **DCAP_0515** | NA | *Drosera capensis* | wide | 8 - DCAP_0515_full_m1_cut.eq.1.pdb | 4 | Fisk_Chem 341L |
| **DCAP_2122** | NA | *Drosera capensis* | wide | 9 - DCAP_2122_full_m1_cut.eq.1.pdb | 4 | Fisk_Chem 341L |
| **Papain** | PAPA1_CARPA | *Carica papaya* | NA | PAPA1_CARPA.pdb | 5 | Both |
